# Optimal enzyme profiles in unbranched metabolic pathways

**DOI:** 10.1101/2023.06.30.547243

**Authors:** Elad Noor, Wolfram Liebermeister

## Abstract

How to optimize the allocation of enzymes in metabolic pathways has been a topic of study for many decades. Although the general problem is complex and non-linear, we have previously shown that it can be solved by convex optimization. In this paper, we focus on unbranched metabolic pathways with simplified enzymatic rate laws and derive analytic solutions to the optimization problem. We revisit existing solutions based on the limit of mass-action rate laws and present new solutions for other rate laws. Furthermore, we revisit a known relationship between flux control coefficients and enzyme abundances in optimal metabolic states. We generalize this relationship to models with density constrains on enzymes and metabolites, and present a new local relationship between optimal reaction elasticities and enzyme amounts. Finally, we apply our theory to derive simple kinetics-based formulae for protein allocation during bacterial growth.

## 1 Introduction

The idea that living beings show optimal shapes or behavior has a very long history. A process like evolution, which combines random mutations with a selection for favorable properties, could potentially lead to optimization, but the question of *if* and/or *when* should we expect living beings to function optimally has been widely debated and is far from solved. In practice, it can be useful to invoke optimality principles to seek insights and design principles that might be relevant also in naturally evolved systems [1]. Specifically, cell metabolism has often been studied using this approach [2, 3], thanks to the powerful mathematical models that we have to describe it. But although natural selection has been the main inspiration for this study, the evolutionary aspects of pathway optimization are not discussed here, and are rather left for the reader to reflect upon.

Within cells, protein is arguably the most important and central resource, both in terms of contributing to fitness, but also since protein synthesis requires large amounts of energy, metabolic precursors, and ribosomes, and the proteins themselves occupy a significant portion of cellular space. Therefore, a cell should generally save protein wherever it can. This notion, specifically for enzymes, has been mathematically applied in genome-scale metabolic models [4, 5], models of core metabolism [6], and in direct comparisons between pathways [7, 8]. Since the proteome is a limiting resource, its allocation to different sectors (metabolism, translation, etc.) is a topic of high interest. In some works people assumed optimality. Other papers assumed a general rule based on linear growth rate control. Interestingly, even a very simple linear chain model with two reactions, representing metabolism and protein synthesis, and a bound on the total protein budget has been successful in explaining bacterial growth laws and overflow metabolism [9, 10]. However, these bacterial growth law models did not consider enzyme kinetics.

Here, we focus on a special case of this cost/benefit analysis: the efficient use of metabolic enzymes in unbranched pathways operating at steady-state, giving priority to scenarios that can be solved analytically. We explore several types of kinetic rate laws and introduce the idea of bounding the total metabolite concentration (which is required in some cases for meaningful results). To define states of maximal enzyme efficiency, we can consider two equivalent optimality problems: maximizing a production flux at a given enzyme budget or minimizing protein usage at a given required production flux. In both cases, we maximize the production flux per enzyme usage within the given constraints. If the product of the pathway is directly tied to biomass, the overall enzyme efficiency, called “biomass/enzyme efficiency”, can serve as a proxy for cell growth [6]. Furthermore, this optimality problem is also relevant in other contexts, such as metabolic engineering of synthetic pathways using a set of existing and/or new-to-nature enzymes with known kinetic parameters [11, 12].

Another perspective often used to analyze metabolic systems is through their control, e.g. the effect of changes in a level of an enzyme on pathway flux [13]. In optimal metabolic states with a bound on the sum of enzyme levels, each enzyme effectively carries an opportunity cost. This cost must be balanced by a marginal benefit, given by the flux control coefficient – as defined in Metabolic Control Analysis (MCA) [14, 15, 13]). Hence, for systems in optimal states, there are simple relationships between enzyme abundance and flux control [16, 17, 18]. We will recapitulate these results below, generalise them to models with a density constraint on enzyme and metabolite levels and illustrate them using some of our analytic solutions. In addition, we present a simple rule that links optimal enzyme investments around a given metabolite to the reaction elasticities of this metabolite.

Finally, we show how the analytic solutions derived here might be useful for modeling high-level phenomena such as the Monod curve (i.e. the relationship between the concentration of a limiting nutrient and the growth rate of bacteria [19]). We use this model to demonstrate how each kinetic parameter should affect the growth rate under different conditions.

In summary, this paper revisits the question of optimal enzyme allocation and adds to previous results. We focus on unbranched metabolic pathways, extend the optimality problem, discuss new optimality conditions, and present analytic solutions that directly show how different factors determine optimal enzyme levels and fluxes. We discuss general principles, in particular how optimal enzyme levels reflect flux control and local reaction elasticities, and use our theory to derive formulae for kinetics-based bacterial growth.

## 2 Results

### 2.1 Optimal states of unbranched pathways

One of the first attempts at analytically solving the enzyme allocation problem was published by Waley [20], who studied short pathways of 2-3 reactions while assuming the concentrations of metabolites (which were denoted *linking intermediates*) are much below the enzymes’ *K*_M_ values, and therefore affect the flux linearly. Given the total amount of catalytically active protein (bounded by *ε*_tot_), the relative enzyme concentrations should be such that they maximize rate (see Figure 1). Based on these assumptions, one can derive simple formulae for the optimal enzyme levels and maximal pathway flux. Later studies repeated this result and generalized it to linear pathways of any size [21, 17, 18]. Here, we will revisit this general solution and extend it to other types of rate laws beyond the one considered by Waley [20] (which, from now on, we will refer to as *mass-action*).

**Figure 1.**
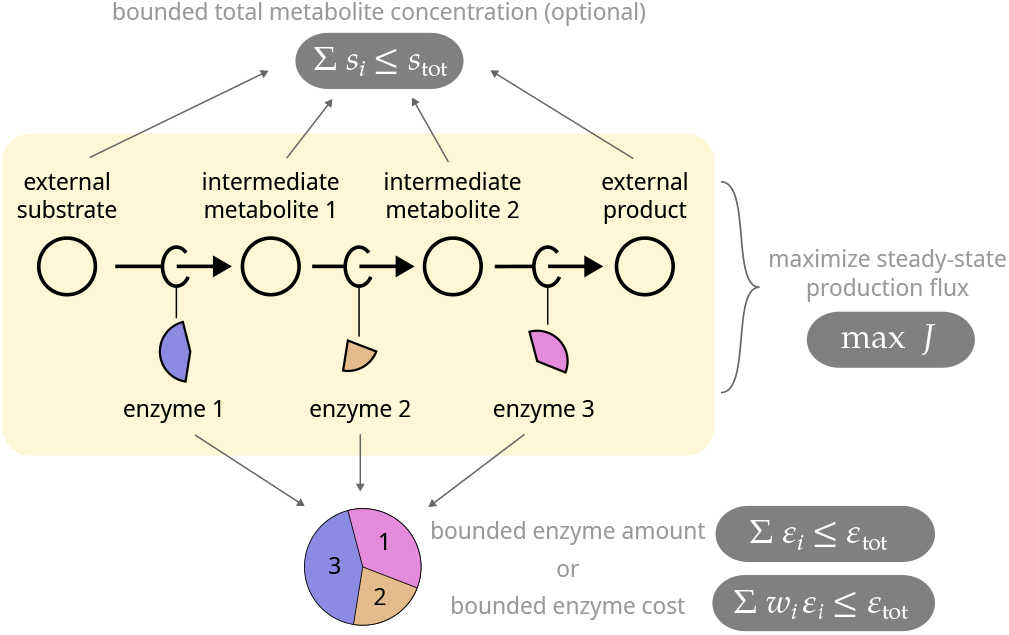
Optimal enzyme levels in unbranched metabolic pathways. In the basic optimality problem in this paper, we consider a chain of reactions and ask how a given protein budget should be spent on metabolic enzymes to achieve a maximal steady-state flux. If all the efficiencies (i.e. the reaction rates per catalyzing enzyme) of all the enzymes were known, the steady-state flux would determine the enzyme levels, and there would be nothing to optimize. Here we assume that the enzyme efficiencies can be adjusted by choosing the metabolite concentrations (not shown), and we search for the optimal metabolite and enzyme profile. The aim is to maximize the flux at a given total enzyme amount (bottom) and possibly under a constraint on metabolite concentrations (top). The flux ratios between reactions are predefined, for example assuming equal fluxes in all the reactions (right).

Consider the following unbranched pathway [22] (Figure 1):

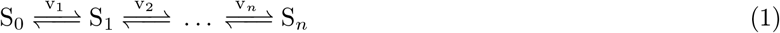

In a kinetic model, each variable *s*_*i*_ represents the concentration of a metabolite *i* and each variable *ε*_*i*_ represents the level (molar concentration or mass concentration) of the enzyme catalyzing reaction *i*. Imagine that the total enzyme level in the pathway is bounded by *ε*_tot_, i.e.

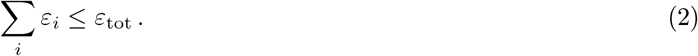

What would be the optimal strategy for distributing this resource between the reactions in order to maximize the steady-state flux? To answer this question, we need to know how the rate of each reaction depends on the levels of enzyme, substrate, product, as well as kinetic parameters. This is described by rate laws. For some rate laws, we can solve the optimization problem and obtain an analytic solution that describes exactly how much of each enzyme should be allocated. Below, we will also consider a variant of this problem with an extra bound on the sum of metabolite levels or with fixed initial and final metabolites (*s*_0_ and *s*_*n*_).

Since single analytic solutions are rare but instructive, we explore them in this article. We consider four different rate laws (summarized in Figure 2): the general Haldane rate law (saturable and reversible) which has no analytic solution, and three solvable approximations derived from it. As explained above, the *enzyme levels ε*_*i*_ may either refer to molar concentrations or mass concentrations (depending on the modeler’s preference). In the case of molar concentrations, the *k*^cat^ values are catalytic constant (e.g. in units of 1/s). In the case of mass concentrations, the *k*^cat^ values are specific activities (e.g. in units of μmol *×* min^*−*1^ *×* mg^*−*1^).

**Figure 2.**
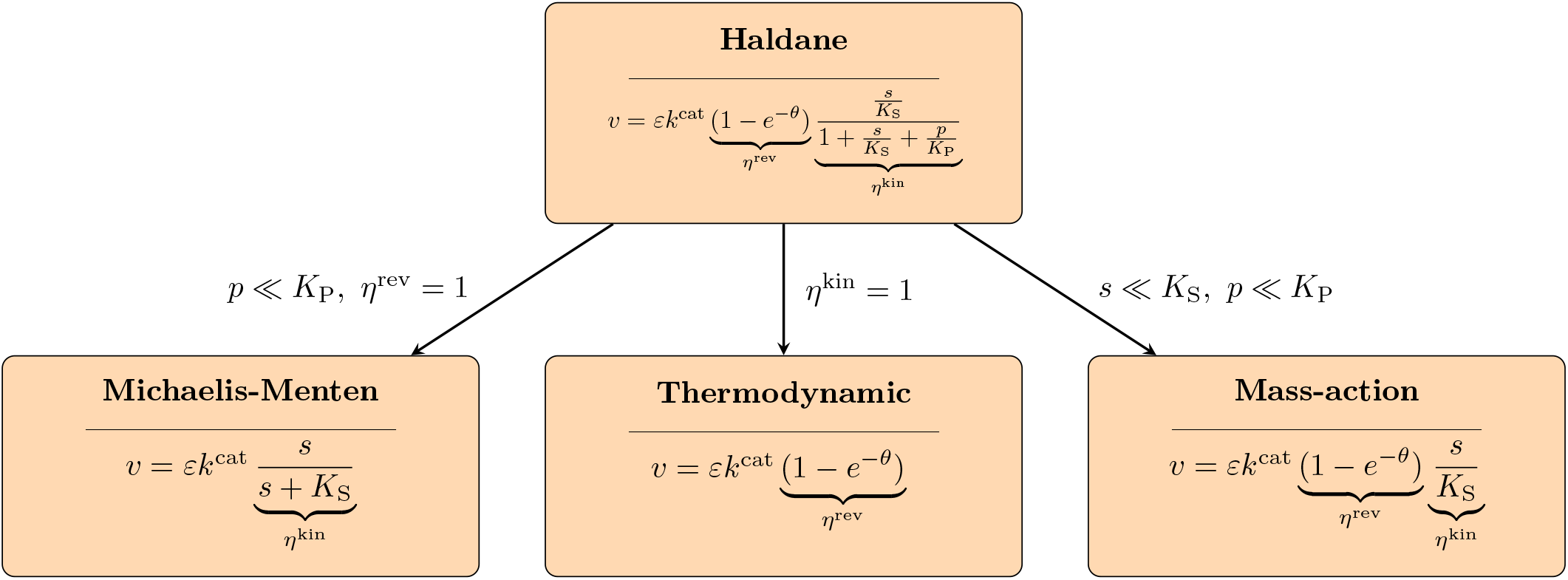
The Haldane rate law and some simplified rate laws. Simplified rate laws are obtained as limiting cases by setting efficiency terms *η*^rev^ or *η*^kin^ to 1 or another constant value (see Figure 3) or by assuming that reactant concentrations are small (in the case of the mass-action rate law). For the enzyme optimality problem in unbranched pathways, we do not know of any analytic solutions for the Haldane rate law. We report here solutions for the other, simplified rate laws (where the solution for the thermodynamic rate law contains an unknown auxiliary parameter).

- **Reversible saturable rate law (“Haldane”)** As a general rate law for a reaction S ↔ P, we consider the reversible saturable rate law

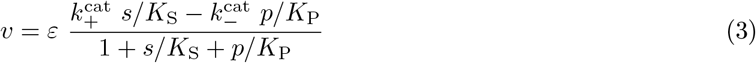

with Michaelis-Mention constants *K*_S_ and *K*_P_, which can be factorized into

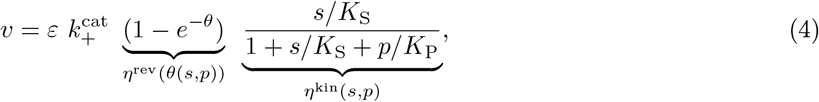

with the thermodynamic driving force *θ*(*s, p*) = ln(*K*^eq^ *s/p*), the thermodynamic efficiency *η*^rev^ and the kinetic efficiency *η*^kin^ (see Figure 3). Note that the driving force can also be defined as *−*Δ*G*^*′*^ which is equal to *RT* ln(*K*^eq^ *s/p*), but here we drop the gas constant *R* and temperature *T* to have a unitless (positive) variable. The two efficiency terms can only assume values between 0 and 1. This factorized formulation of the Haldane rate law is equivalent to the one in Eq. (3), based on the constraint that 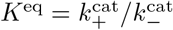, which is commonly known as the *Haldane relationship* (see [23] for more details). This rate law is the most realistic one that we discuss in this work, and deriving it requires only a few assumptions. However, it is also the most mathematically complex and therefore most questions we raise below do not have analytic solutions. So, in addition, we consider three simplified rate laws as limiting cases.
- **Mass-action rate law** A very common approximation for enzymatic rate laws (the one also made by Waley [20]) is based on the limit of low metabolite concentrations (*s ≪K*_S_ and *p ≪ K*_P_). In this case, the concentration-dependent terms in the denominator (i.e., *s/K*_S_ + *p/K*_P_) can be neglected, and we get:

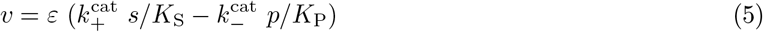

There are many equivalent ways to write down this rate law. For instance, we can apply the same approximation to the factorized form in Eq. (4) to get 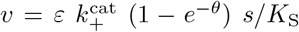, and we can further replace *θ* with its explicit definition based on reactant concentrations and write 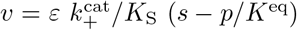. Another common form for this rate law is based on the first-order rate constants 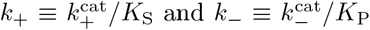. Using them in Eq. (5) looks like this: *v* = *ε* (*k*_+_*s − k*_*−*_*p*). As the rate law resembles mass-action kinetics for non-enzymatic reactions, we will refer to this as the “mass-action” rate law although here the enzyme level appears as a prefactor. Throughout this paper we will switch between these four different notations based on convenience.
- **Thermodynamic rate law** If the kinetic efficiency *η*^kin^ is approximated by 1 (e.g. in the limit *s ≫ K*_S_ and *p ≪ K*_P_), we obtain:

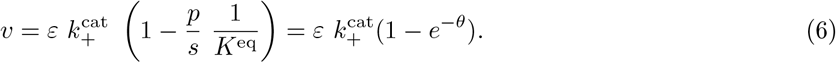

We will also consider a special case of this rate law where *θ* →0 and therefore *p/s* ≈*K*^eq^, i.e. the reaction is close-to-equilibrium. In this case the rate law becomes 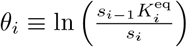.
- **Irreversible saturable rate law (“Michaelis-Menten kinetics”)** We next assume that both *p* ≪*s K*^eq^ (which means that *θ* → ∞and therefore the thermodynamic efficiency *η*^rev^ can be approximated by 1) and also *p ≪ K*_P_ (so we can drop *p/K*_P_ from the denominator in *η*^kin^). In this case, we obtain the MichaelisMenten rate law

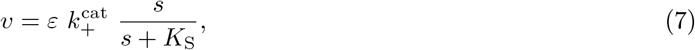

which depends on the substrate, but not on the product concentration. Originally, Michaelis and Menten [24] developed this irreversible rate law by assuming that the rate of enzyme-substrate binding is very fast compared to catalysis, and that the catalytic step is irreversible. The assumptions made here lead to the same result but are less stringent.

**Figure 3.**
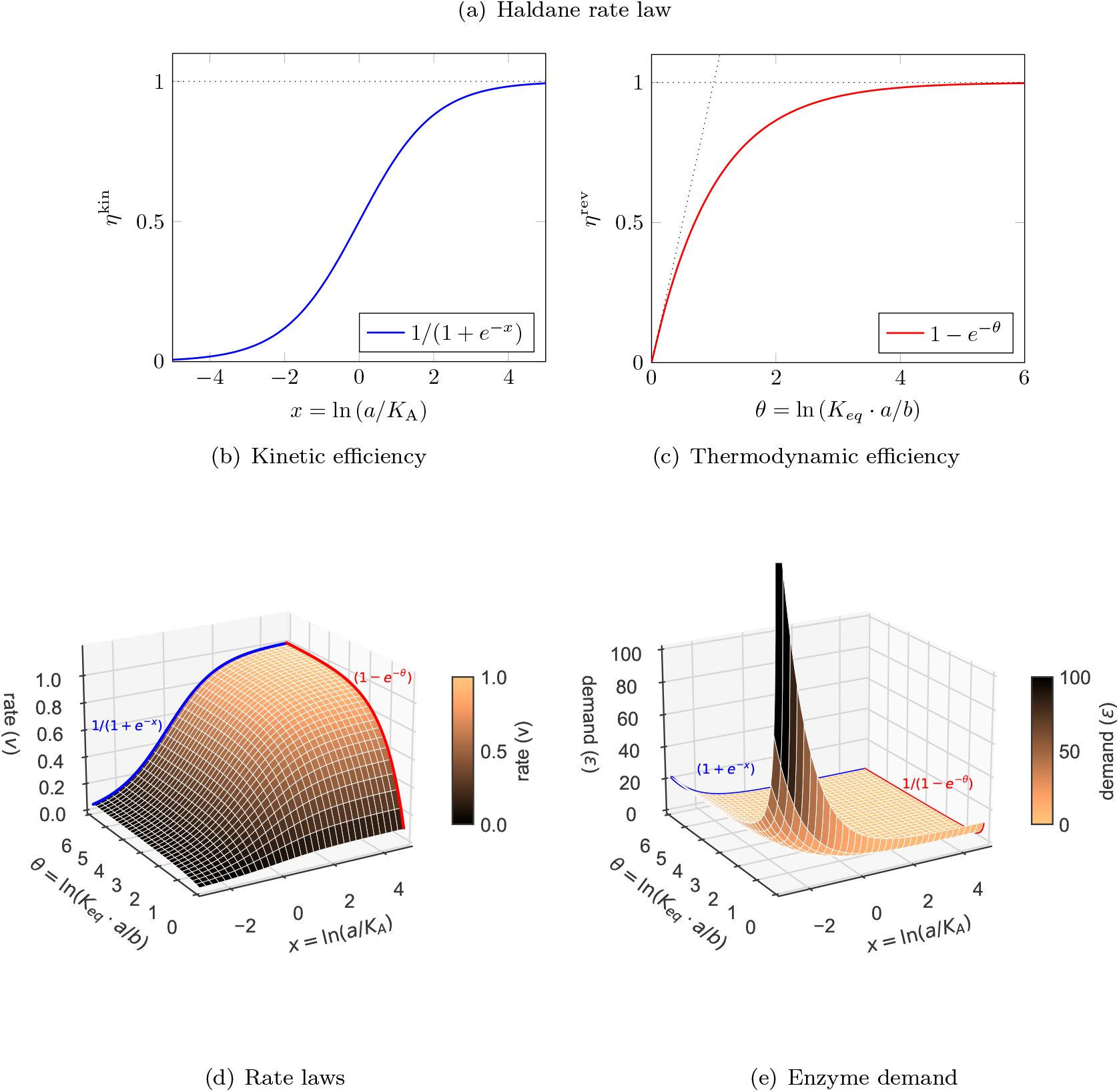
The Haldane rate law and some approximations with constant efficiency terms. (a) The Haldane rate law. In the factorized form (right), thermodynamic and saturation effects are described by separate efficiency factors. (b) The kinetic efficiency as a function of the product of the substrate (in log-scale), assuming *b* ≪ *K*_B_. (c) The thermodynamic efficiency as a function of the driving force *θ*. (d) A surface plot showing the rate *v* as a function of *θ* and the substrate concentration (in log-scale, relative to the *K*_A_). The parameters are *K*^eq^ = *K*_A_ = *k*^cat^ = *ε* = 1 and *K*_B_ = 10. (e) A surface plot showing the enzyme demand for a given rate (*v* = 1). All kinetic parameters are the same as in (d).

In the approximations, instead of setting the thermodynamic or saturation efficiencies to their maximal value of 1, we may also approximate them by constant numbers smaller than 1; we obtain exactly the same rate laws, but instead of the *k*^cat^ value we obtain a smaller apparent value in the approximated rate law.

### 2.2 Analytic formulae for unbranched metabolic pathways

How can we characterize metabolic states by using analytic formulae? Here we consider chains of uni-uni reactions. For such an unbranched pathway with a given type of rate laws (e.g. mass-action or Michaelis-Menten rate laws), we are interested in formulae for a number of quantities:

1. **Metabolic steady state** Given the enzyme levels, external metabolite concentrations, and kinetic constants, we can directly compute the stationary fluxes and internal metabolite concentrations. For general metabolic networks, no explicit formulae are known, but for unbranched pathways with some simplified rate laws, explicit formulae exist. Incidentally, this also shows that in these models the steady state concentrations are unique.
2. **Stability of steady state** If the Jacobian matrix in a steady state has positive eigenvalues, the state is asymptotically unstable and is not able to persist under (inevitable) chemical noise in the cell. Stability is a prerequisite for metabolic control coefficients being defined. A sufficient (but not necessary) condition for stable steady states in unbranched metabolic pathways is given in Appendix F.8.
3. **Metabolic control** The response coefficients are defined as the derivatives between steady-state concentrations or fluxes and model parameters (e.g. the enzyme levels). If two model parameters act (exclusively) on the same reaction, all their response coefficients will be the same (except for a proportional scaling). Taking this into account, the control coefficients describe the same type of derivatives, but for a set of hypothetical, reaction-specific parameters. In practice, if reaction rates are proportional to enzyme levels and if each reaction is catalyzed by a single specific enzyme, we can write the enzyme response coefficients as 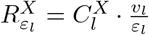. Importantly, throughout this paper we use elasticities, response coefficients, and control coefficients in their unscaled form, for instance unscaled enzyme response coefficients 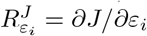 instead of the common scaled form 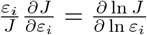. The metabolic control coefficients can be computed from the stoichiometric matrix and the elasticity matrix, and they satisfy summation and connectivity theorems. They can be computed in two ways: if an analytic formula for the metabolic steady state is known (as is the case in some unbranched pathway models studied below), we may differentiate symbolically by the enzyme levels; otherwise, we may compute the elasticity coefficient matrices by differentiating the rate laws, and then compute the control coefficient matrices from them using a known formula; however, since this formula involves a matrix inversion, writing this down as an analytic formulae may be extremely complicated, and control coefficients are usually computed numerically. Another way to compute control coefficients, which follows from the enzyme-control rule and works only in optimal states, is described below.
4. **Optimal metabolic states** In our basic metabolic optimality problem, we define optimal states as states in which a given enzyme budget (fixed sum of enzyme levels) is allocated to maximize a production flux. Kinetic constants and external metabolites are given, and we compute the optimal metabolite profile, the optimal enzyme profile, and the optimal flux. If the flux distribution is known (e.g. a steady flux in an unbranched pathway) and can only increase or decrease proportionally, this problem is equivalent to the ECM problem of minimizing the enzyme demand at a given (unit) flux. This convex problem can be solved numerically, but analytic solutions were known only for very few cases. Below we present some new analytic solutions.

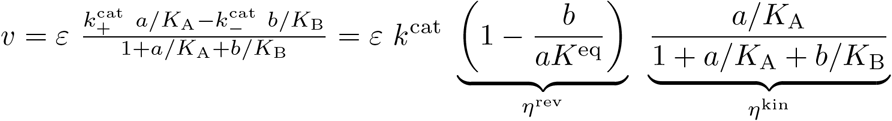

Formulae for optimal enzyme levels and the optimal achievable flux are shown below, in Table 1. We also consider a related problem, maximizing the flux under a constraint on the total enzyme plus metabolite amount.

**Table 1:**
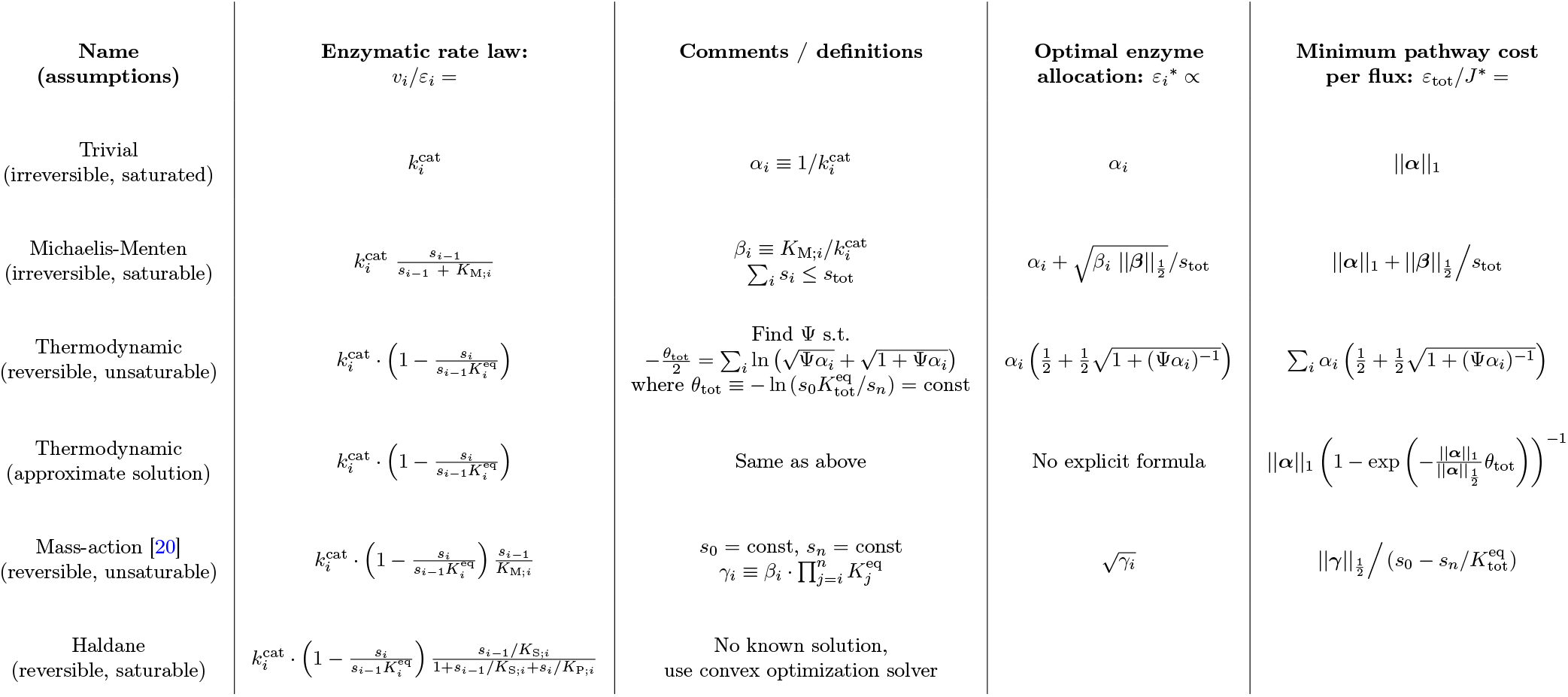
A summary of all rate laws considered in this paper, along with their solutions for optimal enzyme allocation and the resulting minimum pathway cost per flux. The rate laws are roughly ordered by increasing level of complexity. The “trivial” rate law does not have its own section in the text, but is shown here as the limit of the Michaelis-Menten rate law when *s*_*i−*1_ *≫ K*_M;*i*_, as well as the thermodynamic rate law when 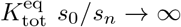. Although the “mass-action” rate law appears in this table in a modified form (for the purpose of using the same set of kinetic parameters as the other rate laws), it is equivalent to the form used in previous studies [20, 16, 17]. In order to calculate the absolute optimal enzyme levels from the relative ones (given in the column titled *Optimal enzyme allocation*), one can use 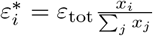 where *x*_*i*_ are the relative values.
5. **Metabolic control in optimal states** The control coefficients tell us how a metabolic system responds to perturbations (resulting in a new state that is stationary, but probably non-optimal). For optimal states (with a constraint on the sum of enzyme levels), the enzyme-control rule states that enzyme levels and flux control coefficients must be proportional. Since we know (from the summation theorem) that the control coefficients must sum to 1, we can conclude: whenever there is an analytic formula for optimal enzyme levels, we obtain a formula for the control coefficients.

### 2.3 Optimal metabolic states: analytic solutions

What are the general principles behind optimal enzyme allocation? One important principle, valid in optimal metabolic states, has been shown for pathways with mass-action rate laws [21] and later been confirmed for general rate laws [17]: in optimal states, where the metabolic flux has been maximized at a fixed enzyme budget, enzyme levels and flux control coefficients must be proportional^1^:

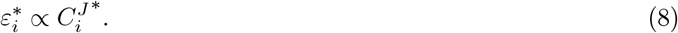

Here the star ^*∗*^ denotes variables in the optimal state. Together with the summation theorem for flux control coefficients [14, 25], 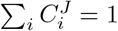, we obtain the conversion formulae

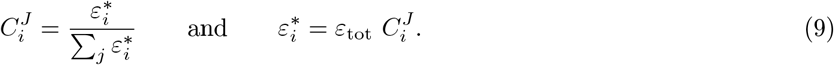

The enzyme-control rule (8) provides a condition for metabolic states, independent of the type of rate laws considered. Importantly, it holds only in states of maximal flux, given a fixed total enzyme amount and no other constraints. More realistic models employ also bounds on metabolite levels [22, 5, 26] for different reasons. First, metabolite levels are bounded in real cells: while metabolite molecules may be small, their concentration is high, and they contribute much more than macromolecules to osmotic pressure. Second, as we will see below, some models without metabolite bounds lead to paradoxical results. We will therefore present a generalized version of the enzyme-control rule that takes metabolite bounds into account.

But what are the general shapes of optimal enzyme profiles, i.e. how do enzyme levels vary across the network? And on what factors (kinetic, thermodynamic, or cost factors) does this depend? To answer this, the enzyme-control rule alone would not be enough (because no kinetic details appear in the rule). Also numerical studies would not be enough (because they apply to single cases and yield no general laws). Hence, to study this, it would be good to consider analytic solutions. Unfortunately, analytic solutions are not known for general metabolic models, but solutions exist for unbranched pathways with simple rate laws (mass-action rate law and saturable kinetics.

We now present analytic formulae for optimal metabolic states with different types of rate laws. We discuss the rate laws in increasing order of difficulty. An overview of all analytic solutions is given in Table 1.

#### 2.3.1 Michaelis-Menten rate law

We first consider the Michaelis-Menten rate law (i.e. irreversible reactions with simple saturation kinetics):

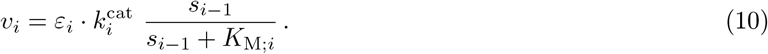

In an unbranched pathway at steady state, all rates must be equal. To describe them, we introduce a new variable *J* called the pathway flux, and require that *∀i v*_*i*_ = *J*. Now we can use Eq. (10) to find a relationship between substrate and enzyme levels:

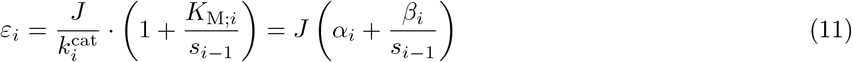

where we defined 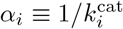 and 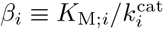 for convenience.

Combining this equation with an upper bound on total enzyme from Eq. (2) we get that:

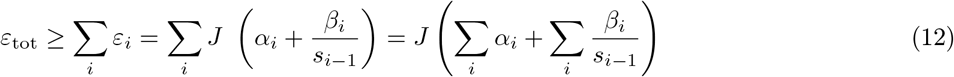

and by rearranging we obtain:

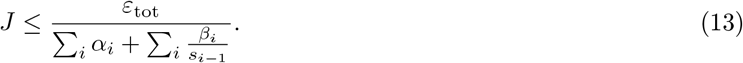

Maximizing *J* would mean that we reach the upper bound, and therefore we can also treat this as an equality. Since *ε*_tot_ is constant and the only free variables are the metabolite concentrations, the maximal flux is reached when the denominator on the right-hand side is minimized. The problem is that it is a monotonically decreasing function in *s*_*i*_ (for each *i*) and since metabolite concentrations are unbounded, the optimum would be at *s*_*i*_ → ∞ which, formally, is not a mathematically defined optimum state. But even if we consider this as a limit, the resulting state would not be controllable, but instead very fragile against any changes of enzyme levels (see Appendix C.1). In reality, of course, the range of physiological osmotic pressures does impose some constraint on the concentrations of small molecules. As a proxy for this effect, we can add another constraint to bound the sum of all metabolite concentrations, Σ _*i*_ *s*_*i*_ *s*_tot_. Now, one can show (see Appendix F.1) that the optimal allocation of enzymes will obey:

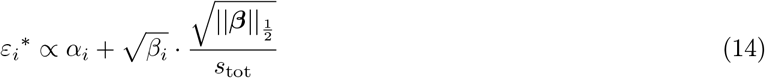

where ***β*** is the vector of all *β*_*i*_ and 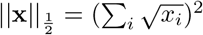 is the 1_1/2_ norm (i.e., 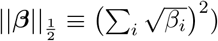). In this case, maximal flux would be:

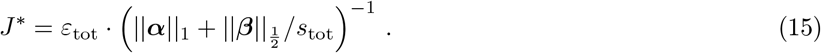

Note that the solution looks essentially the same even if we constrain the first metabolite (*s*_0_) to have a fixed concentration (see Appendix F.1.4). We will revisit this case in section 2.5 in the context of a course-grained model of a growing cell.

If the metabolite density constraint is not very tight (i.e. *s*_tot_ is large enough), we can ignore the second term in Eq. (14), which would be equivalent to assuming all enzymes are substrate-saturated. In this case, the optimal allocation of enzymes will be proportional to *α*_*i*_ (or inversely proportional to 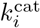) and therefore:

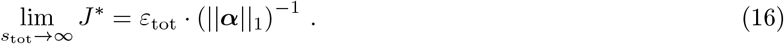

Interestingly, | |***α***| | _1_, which is equal to the sum of *k*^cat^ reciprocals, is in fact the inverse of the *Pathway Specific Activity* (PSA) as originally defined by Bar-Even et al. [11]. Indeed, the idealized scenario considered in that study (where all enzymes were irreversible and saturated) provides an upper bound on the maximal flux achievable. Interestingly, if we see the 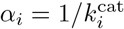 as minimal enzymatic turnover times and *α*_PW_ = *ε*_tot_*/J* is the turnover time of the pathway, we can simply rephrase Eq. (16) by saying that, like in the case of the PSA, turnover times along a pathway are additive. In fact, this holds more generally for any rate law *v*_*i*_ = *e*_*i*_ *k*_*i*_(**s**), if the enzyme turnover times are defined as *τ*_*i*_ = 1*/k*_*i*_, and resembles the fact that electric resistances, in a series of resistors, are additive.

One reason that the irreversible Michaelis-Menten model is unrealistic is that it does not capture reactions that are close to equilibrium and therefore suffer from a counter-productive reverse flux. This is yet another reason why it might be impossible for some metabolites to reach very high concentrations: they might be products of unfavorable reactions. In most metabolic networks, about half of the reactions are reversible and therefore it would be more realistic to use a reversible rate law like such as Haldane’s. However, using the Haldane equation would typically create a system of equations for which no analytic solution is known. So, first, we will consider rate laws that account for the reverse flux but, for simplicity, ignore substrate saturation.

#### 2.3.2 Thermodynamic rate law

Irreversible rate laws like the Michaelis-Menten kinetics depend only on the substrate concentration; in the formula there is no product-dependent reverse term that decreases the total rate or could make it become negative. However, according to thermodynamics, such laws can only be approximations: thermodynamically feasible rate laws must contain a reverse term and must depend on the thermodynamic imbalance of substrate and product concentrations expressed by the thermodynamic driving force. Some rate laws can even be written as functions of the thermodynamic force alone. Here we describe such a “thermodynamic” rate law, where *v* is proportional to the thermodynamic efficiency 1 *−* e^*−θ*^, while the kinetic efficiency *η*^kin^ is assumed to be constant:

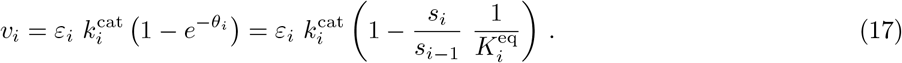

This type of kinetics approximates cases where all reactions are saturated (*s*_*i*_ ≫*K*_M;*i*_), and therefore the thermodynamic efficiency *η*^kin^ in Eq. (4) becomes 1. The parameters here are the turnover numbers 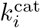 and the equilibrium constants 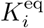. However, as we will soon learn, the individual 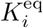 do not change the result, and only the overall equilibrium constant 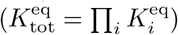 matters.

As before, we can define the steady-state pathway flux *J* and apply an upper bound on the total enzyme to get:

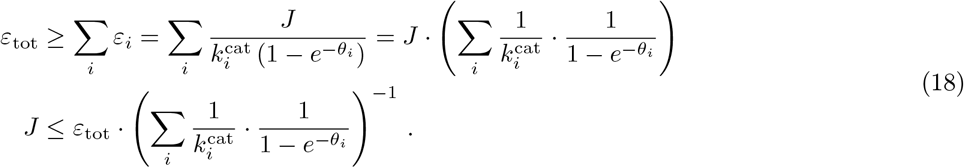

Unfortunately, there is no closed-form analytic solution for the maximal rate of this thermodynamic rate law. But we do have something very close which requires rather simple computations. First, we need to invert the functional relationship between the overall driving force (*θ*_tot_) and an auxiliary variable Ψ which is defined by the inverse function of:

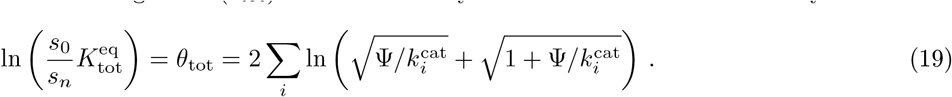

The expression on the right is analytic and strictly increasing in the range Ψ ∈ [0, ∞), so there is a unique solution which can be found by simple numerical methods. Then, we can use that value to directly calculate the optimal driving forces, enzyme levels and pathway flux:

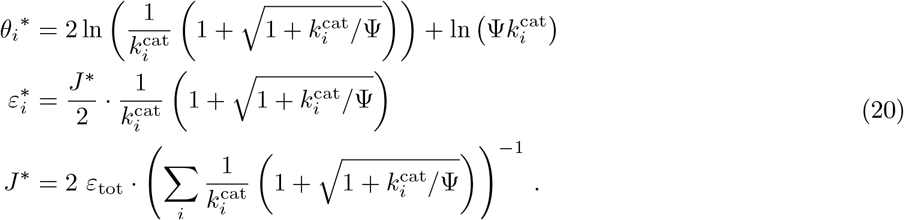

The full derivation of this solution can be found in Appendix F.2.2. An example of what the relationship between the driving force and the optimal flux looks like for a pathway with two enzymes is illustrated in Figure 4(a) and 4(b). Furthermore, a comparison between our exact formula presented here and a numerical solution based on convex optimization [8] shows no difference at all.

**Figure 4.**
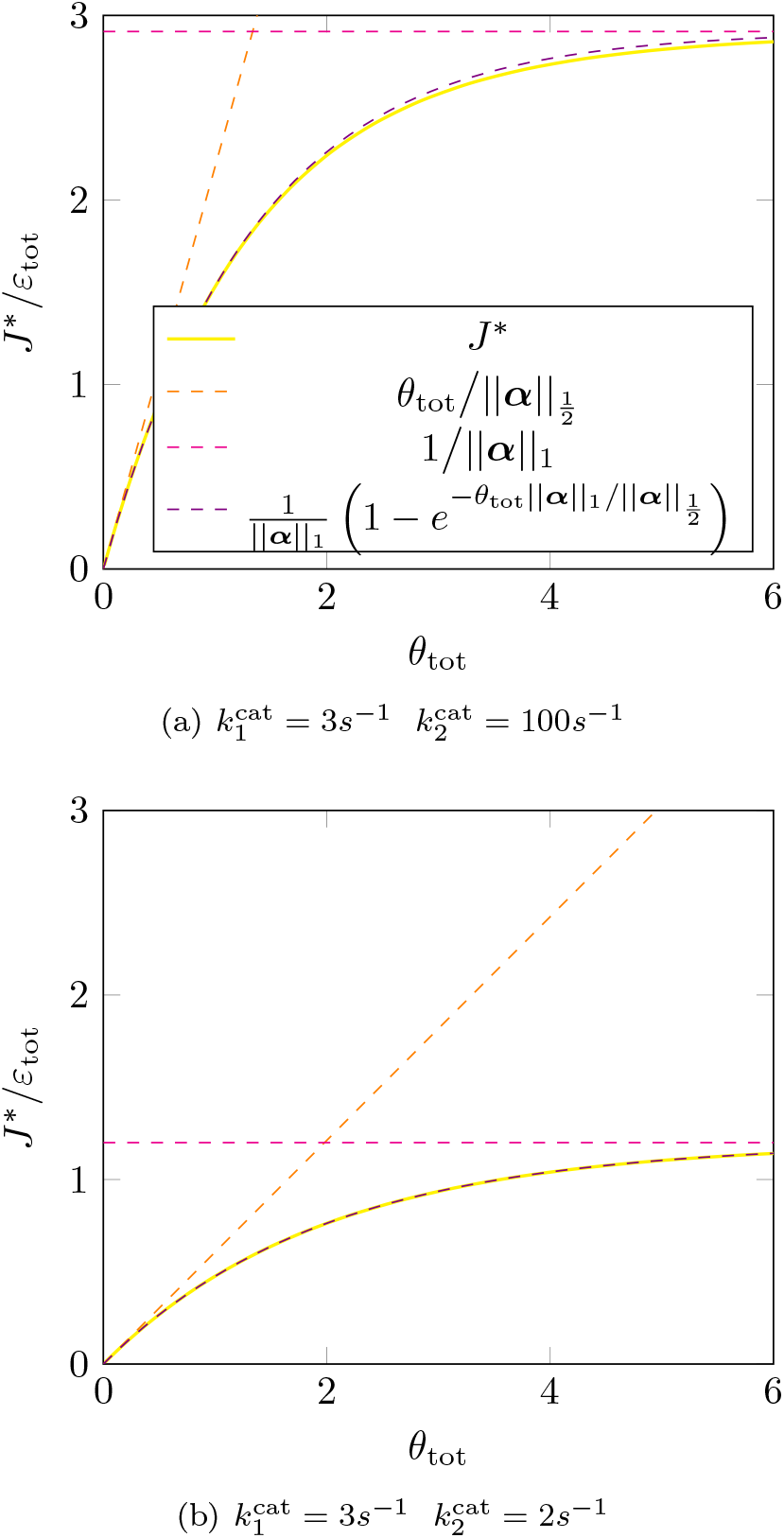
The relationship between the maximal flux per enzyme and overall driving force for the thermodynamic rate law. Even though we do not have a closed-form solution for *J* ^*∗*^ as a function of *θ*_tot_, we can still plot *J* ^*∗*^(Ψ) against *θ*_tot_(Ψ) for varying values of Ψ using Eq. (19) and Eq. (20). Here, we show this relationship for a pathway with 2 steps (yellow). The parameters are: 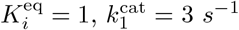, and the *k*^cat^ of the second enzyme is either (a) 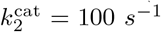 or (b) 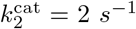. The orange dashed line represents the approximations for very small driving forces based on Eq. (23), and similarly in pink for very large driving forces – Eq. (21). The approximation (dashed purple line) based on Eq. (24) is based on effective parameters chosen to match the slope at *θ*_tot_ = 0 and the limiting maximal flux at *θ*_tot_ *→ ∞* .

These formulae cannot be directly evaluated because of the unknown parameter Ψ. To obtain solutions that do not depend on this parameter, we now consider two limiting cases in which Ψ is either infinitely high (very high driving force) or infinitely low (very low driving force).

When the driving forces are very high (i.e. *θ*_tot_ → ∞), the solution for Ψ in Eq. (19) will approach infinity and the optimal flux becomes

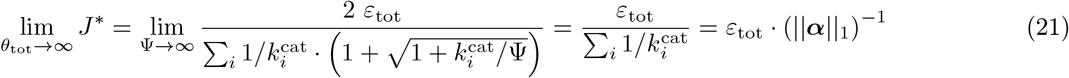

with parameters 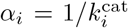 defined as before. This solution indeed makes sense as it is equivalent to the fully saturated limit in the Michaelis-Menten case (see Section 2.3.1, Eq. (16)).

However, when the driving force in the pathway is limited (Ψ has a finite positive value), this driving force will be split between reactions according to their 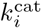 values: enzymes with a higher 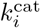 value will have to pay a higher penalty due to their driving force being closer to 0. On the other hand, slow enzymes will obtain more driving force, which will help them by having a smaller fraction of reverse flux. Notably, the distribution does *not* depend on the reaction equilibrium constants (only on the overall 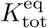). Perhaps this is not that surprising if we consider the fact that the concentrations intermediate substrates and products are unconstrained and therefore we have enough degrees of freedom to adjust to any value of 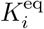 as long as the total driving force stays the same.

One can also consider the other extreme where all reactions are close to equilibrium, which means that *θ*_tot_ is close to 0 (and therefore also each *θ*_*i*_ *→* 0). In this case, we can approximate Eq. (18) by:

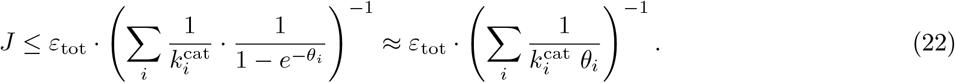

Maximizing *J* under the constraint Σ_*i*_ *θ*_*i*_ = *θ*_tot_ yields the solution (see the full derivation in appendix F.2.3):

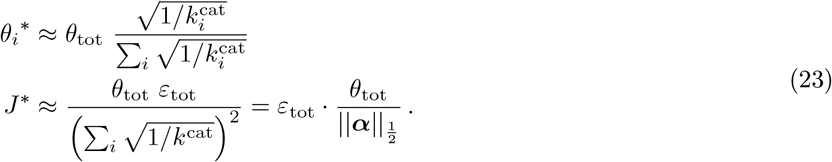

Using the two limiting cases (*θ*_tot_ → ∞and *θ*_tot_ ≈ 0), we can approximate the solution from Eq. (20) by a much simpler formula which has the same shape as the thermodynamic rate law for a single reaction:

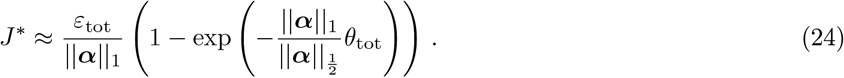

Figures 4 compares between the precise and approximate solution for two example cases (a more detailed comparison for metabolic pathways of different lengths and parameter choices is shown in Appendix F.3). One can appreciate that the two curves are nearly indistinguishable, illustrating the good quality of the approximation.

#### 2.3.3 Mass-action rate law

Next, we consider the same unbranched pathway but assuming mass-action rate laws:

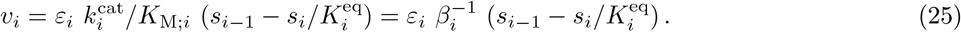

As before, we defined 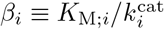 Note that instead of Eq. (5) we use a form of the mass-action rate law that does not require the turnover rate and *K*_M_ of the product (and instead uses 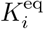), to avoid confusing indexation of forward and backward parameters.

Like with the previous rate laws, we define the pathway flux *J* and apply an upper bound on the total enzyme levels:

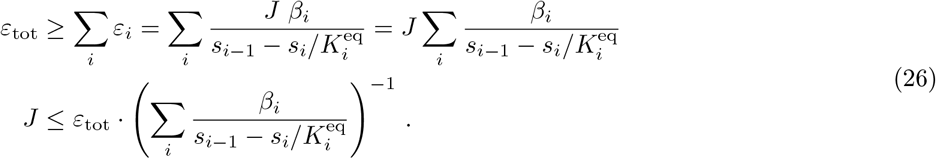

Again, we maximize the flux at a constrained total enzyme level. Optimization with Lagrange multipliers yields the following expressions for the optimal individual enzyme levels (*ε*_*j*_^*∗*^) and the maximal flux (_*J*_ ^*∗*^) (see full derivation in Appendix F.4):

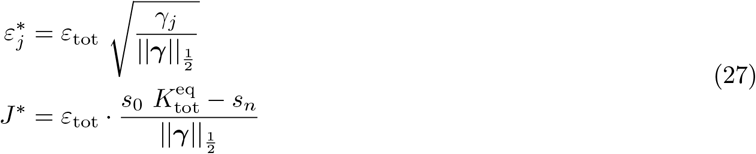

where *γ*_*j*_ is defined as

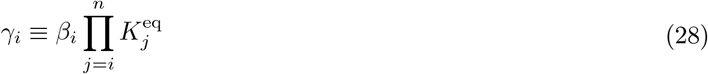

Of course, the exact value of this maximum depends on all the different system parameters. However, it is interesting to consider a naïve assumption where all the *γ*_*j*_ parameters are identical. In such a case, the flux in the pathway would decrease quadratically with the number of steps [13]. Of course, we know that in real metabolic pathways the equilibrium constants are typically not close to 1, and therefore this approximation might not have many applications in biology.

What would happen if a mutation improved the catalytic rate of only one of the enzymes *ε*_*i*_ by a factor of *a* (i.e. *β*_*i*_ decreases by a factor of *a*, but the equilibrium constant 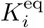 remains the same)? In this case, *γ*_*i*_ would be divided by *a* but the optimal enzyme concentration for this reaction *ε*_*i*_^*∗*^ would only decrease by a factor of 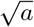. This saving would then be distributed proportionally among all the *n* enzymes and would contribute to an increase in the pathway flux *J*. On the flip side, if for some reason the activity of one enzyme is decreased by a multiplicative factor *b*, it would need to “pay” (increase the enzyme’s concentration) only by a factor 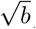, and this increase would be “funded” by all of the enzymes together in order to keep the same *ε*_tot_, thus lowering *J*.

#### 2.3.4 Haldane rate law

So far we analyzed three cases where all enzymes could be described by the same kinetic rate law: mass-action, Michaelis-Menten, or thermodynamic. Although these rate laws can reliably describe some enzymes in specific conditions, it is very unlikely that such an approximation would apply to *all* the reactions in a single pathway (except for the trivial case of a 1-reaction pathway). A more realistic model would allow all reactions to be reversible and saturable. Here, we will analyze such a case based on the factorized rate law with one substrate and one product. Note that although it is equivalent to Eq. (3), we prefer this formulation because it is easier to separate thermodynamics from saturation effects.

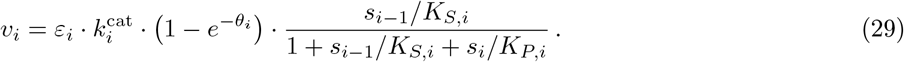

As always, we can assume that all fluxes are equal to *J*, and use the total enzyme budget to get an upper bound:

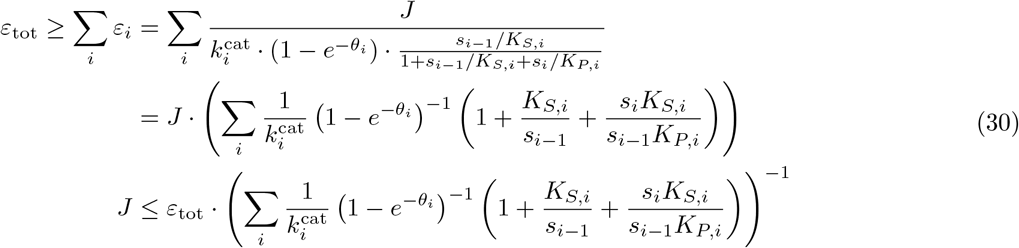

where we can now appreciate how this is a generalization of both Eq. (13) (Michaelis-Menten) and (18) (thermodynamic). We can see that the maximal pathway flux would be realized when the term in parentheses is minimized with respect to the *s*_*i*_, i.e.:

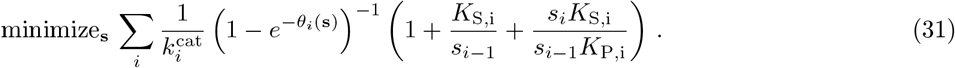

Unfortunately, as we already mentioned before, no analytic solution to this problem is known in the general case.

### 2.4 Optimal metabolic states: insights from Metabolic Control Analysis

#### 2.4.1 Enzyme-control rule and enzyme-elasticity rule

We now continue with some observations about the enzyme-control rule. The enzyme-control rule Eq. (8) states that enzyme levels and flux control coefficients in optimal states are proportional: 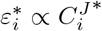. When the aim is to compute control coefficients in optimal states (at a given enzyme budget *ε*_tot_), the enzyme-control rule comes in handy. The summation theorem tells us that the flux control coefficients must sum to 1; therefore, the optimal enzyme levels are given by the flux control coefficients, multiplied by the (predefined) total enzyme level, and we obtain the simple conversion 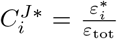 and 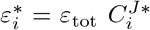. In the basic form of the rule we put a constraint on the enzyme mass, and assume that enzyme levels are mass densities. If the enzyme levels are molar concentrations, and differently weighted in the constraint, the weights can be taken into account by modifying the rule. To what types of optimality problems does the rule apply? If the metabolic system is not an unbranched pathway, but a general network with one target flux, the rule will still hold (where the flux control coefficients refer to this target flux. This also holds for models with metabolite dilution.

However, there are some limitations. Obviously, the rule cannot hold in states in which control coefficients are not defined. This condition may seem unproblematic, but it is actually violated in the pathway with Michaelis-Menten rate laws, because in the limit *s*_*i*_ → ∞ required by optimization, any variations of single enzyme levels would break the steady state. We discuss this in Appendix C.1. One way to avoid this problem is to add a bound on metabolite levels. For example, we may consider a generalized density constraint on enzyme levels and metabolite concentrations

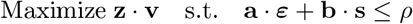

with a linear flux objective **a *ε*** instead of a single target flux, and enzyme weights *a*_*l*_ and metabolite weights *b*_*i*_, and an upper bound *ρ* on the molecule density. Problems with this constraint lead to a generalized form of the enzyme-control rule. For an unbranched pathway with equal weights *a* for enzymes and *b* for metabolites (e.g. molecular masses), we obtain the new enzyme-control rule

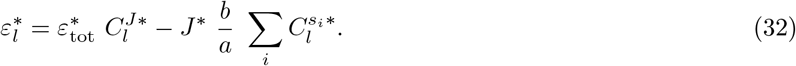

where 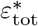 is the (non-fixed) sum of enzyme levels emerging in the optimal state and the symbols 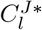 and 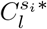 denote, respectively, the flux and control coefficients in the optimal state. For derivation and details, see Appendix C.2.

The original enzyme-control rule has a direct and useful consequence. Since enzyme levels (in optimal states) are proportional to flux control coefficients, they need to satisfy a connectivity theorem. Connectivity theorems relate the elasticities of a given metabolite *i* to the control coefficients of reactions around this metabolite. In the case of flux control coefficients, the right-hand side of the theorem is zero, and we obtain a simple equation for the optimal enzyme levels around a metabolite *i*:

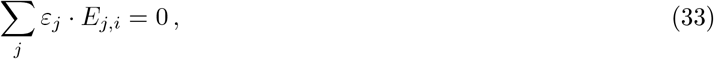

where *E*_*j,i*_ denotes the unscaled elasticity between metabolite *i* and reaction *j*.

We call this formula the enzyme-elasticity rule (see Appendix D). A similar rule for small adaptations of enzyme levels, instead of the enzyme levels themselves, has been shown in [27]. For a linear pathway, the enzyme-elasticity rule yields a simple result: in an optimal state, for each metabolite *i* and its producing reaction *j* = *i −* 1 and its consuming reaction *i*, the ratio of enzyme levels 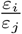 must be equal to the absolute inverse ratio of the elasticities 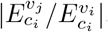. An example of this, with the mass-action rate law, is derived in Appendix D. In contrast to the enzyme-control rule (which yields only one equation for the entire pathway), the enzyme-elasticity rule yields an equation for every internal metabolite, which determines the ratio of the enzyme levels around this metabolite. Together with the known sum of enzyme levels, these rules determine the enzyme profile completely (of course, given all the elasticities in the optimal state).

#### 2.4.2 Analytic formulae for metabolic control

We learned that enzyme levels in optimal states (with a bound on total enzyme, and maximizing the steady-state flux) satisfy two general laws, the enzyme-control rule and the enzyme-elasticity rule. Hence, if we have a formula for the optimal enzyme levels and if the enzyme-control rule applies, we obtain the flux control coefficients. Moreover, the enzyme-elasticity rule (which comes from the connectivity theorem) relates the optimal enzyme levels to elasticities. These formulae can be trusted, but for didactic reasons we computed control coefficients for some of the rate laws analytically (also demonstrating different ways to compute them) and compared them to optimal enzyme levels to check the rules. Here we summarize this briefly; details can be found in the Appendix.

- **Michaelis-Menten rate law** In metabolic pathways with irreversible rate laws, the first reaction has full flux control and therefore a flux control coefficient of 1, while the remaining reactions of no flux control and therefore flux control coefficients of 0. But under an optimization, this changes. All metabolite levels go to infinity and all elasticities go zero. Mathematically, the optimal state doe not exist (because infinite concentrations are not real numbers), but even if we assume that enzymes can be saturated completely, any enzyme variation would break the steady state. This means that the control coefficients are not defined, and so the enzyme-control rule does not apply (see Appendix C.1). To obtain meaningful results, we therefore considered an optimality problem with enzyme and metabolite constraints. In this case, metabolite levels remain finite and the control coefficients remain defined.
- **Thermodynamic rate law** For pathway models with the thermodynamic rate law, we can derive a simple formula for the flux control coefficients that contains only the enzyme levels, the *k*^cat^ values, and the flux. Even if we don’t have an explicit formula for the flux, we can verify that the flux control coefficients satisfy summation and connectivity theorems and that the enzyme-control rule is satisfied. If we do not have a closed formula for the flux as a function of enzyme levels, how can we find a closed formula for the flux control coefficients? Such a formula can be obtained with a trick. From the rate law, we obtain a relation between enzyme levels, external metabolite levels, kinetic constants, and flux:

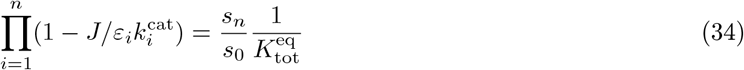

(see Eq. (71) in the Appendix for the derivation). While we cannot solve this for the flux *J* directly, we can obtain the response coefficients 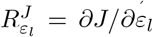 by implicit differentiation. This yields the flux control coefficients (derivation in Appendix F.3.2):

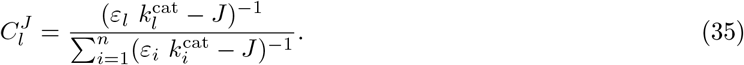

The control coefficients are proportional to 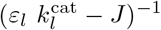 and normalized to a sum of 1 as required by the summation theorem. For didactical reasons, we checked the connectivity theorem (Appendix F.3.1) and the enzyme-control rule (Appendix F.3.4).
- **Mass-action rate law** With the reversible mass-action rate law, we can analytically compute the flux control coefficients and verify that they satisfy summation and connectivity theorems. To verify that the enzyme-control rule is satisfied in the optimal state, we use the explicit formula (26) for the steady-state flux and take derivatives to obtain the flux control coefficients

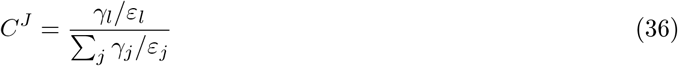

with *γ*_*l*_ defined as above (derivation in Appendix F.4.2). The coefficients sum to 1 as required by the summation theorem. Again, we verified the connectivity theorem (Appendix F.4.4) and the enzyme-control rule (Appendix F.4.5) for didactical reasons.

### 2.5 A cell model with enzyme kinetics and metabolite constraints

As an illustrative example, we apply our formulae to a simple model of a growing cell, describing transport processes, metabolism, and macromolecule synthesis, respectively, by lumped reactions. Obviously, none of the rate laws considered throughout this work can fully capture the dynamics of growing cells. For example, having only one single reaction representing metabolism is a gross oversimplification (and likewise for transport and translation). Nevertheless, we might still be able to draw insights from this model if we make the right assumptions. This approach has been successful in the past: by considering simple cell models and assuming that enzyme efficiencies are constant (i.e. completely independent of growth rate and metabolite concentrations), Basan et al. [10] were able to show how overflow metabolism in *E. coli* corresponds to a proteome-efficient allocation of enzymes. Here we ask how the predictions would change by accounting for enzyme kinetics. Instead of assuming that enzyme efficiencies are constant and given, we consider a model in which protein allocation and metabolite concentrations are always optimized in order to maximize the growth rate.

In our simple cell model (Figure 5) production processes are represented by an unbranched chain of reactions, describing overall transport, overall metabolism, and protein synthesis. The steady-state fluxes are proportional to the cell growth rate and optimized under constraints: an upper bound on the sum of enzyme levels (*ε*_tot_) and another one on metabolite levels (*s*_tot_). For convenience, in this section we will replace the pathway flux *J* ^*∗*^ with the standard symbol for growth rate – *μ*. It should be noted that dilution of the intermediate compounds is ignored even though in reality some of them can have a high total concentration (e.g., the precursors in *S*_2_) and thus a considerable growth dilution rate which affects protein allocation strategies.

**Figure 5.**
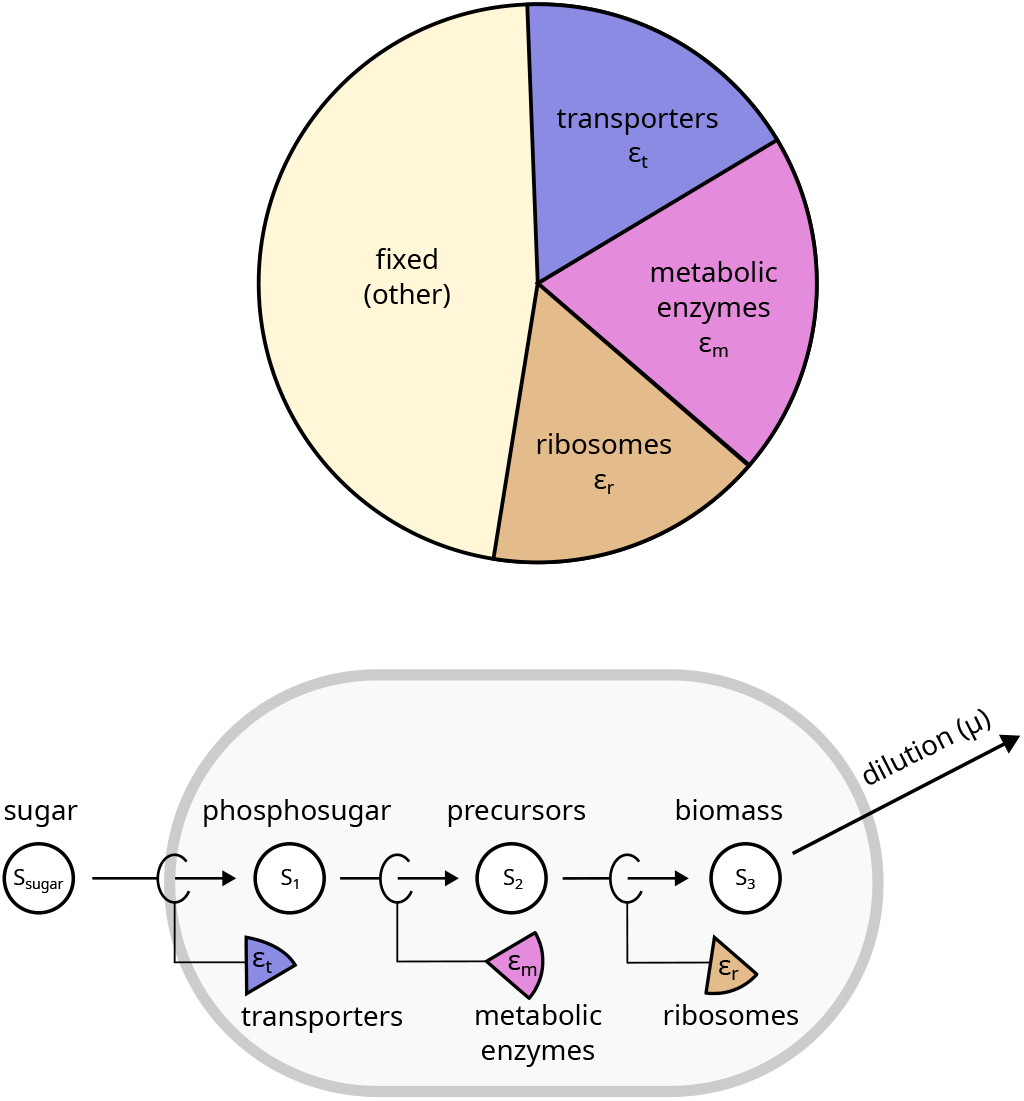
A growing cell described as a chain of reactions. Top: Inspired by the cell model in [9] describing “bacterial growth laws”, we assume a fixed protein budget consisting of a fixed protein fraction and variable fractions for transporters, metabolic enzymes, and ribosomal proteins, but with a fixed sum. The proportions of the variable fractions are chosen to maximize growth. Bottom: The growing cell is described as a chain of reactions, each catalyzed by one of the protein fractions. For a constant biomass production, the three reactions must be in steady state.

For our model, we chose to use the Michaelis-Menten approximation, so that metabolite concentrations (as extra variables) can be adjusted and become part of the optimization problem. We then apply the formulae derived in the previous sections of this paper to find the optimal allocation of enzymes and thereby maximize the growth rate of the cell. Importantly, all calculations are completely based on analytic expressions.

In section 2.3.1, we derived a formula for the optimal allocation and maximal flux in the case where all metabolite levels are free variables (including *s*_0_, which is denoted here by *s*_sugar_). However, since in this cell model *s*_sugar_ represents the concentration of an external nutrient that is not subject to optimization, we would like to treat it as a constant system parameter (and later show how the optimum responds to changes in it). Fortunately, adding this assumption changes the optimal solution only slightly, as described in Appendix F.1.4, and the optimal growth rate as a function of *s*_sugar_ can be written in the following simple form:

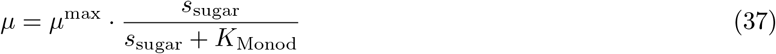

where we define

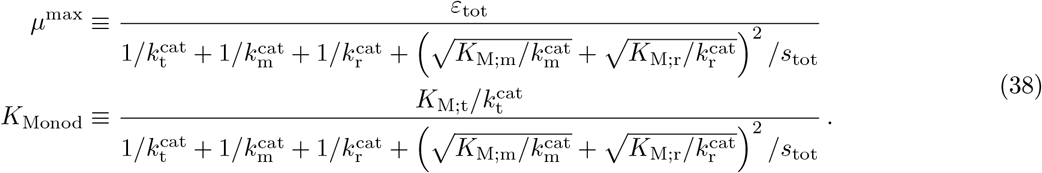

As implied by the symbol for *K*_Monod_, this form corresponds to empirical observations by Monod [19] (and further followed up by others [28]) which stated that growth rate increases with the nutrient concentration until reaching a saturation level where growth is fastest. Interestingly, the value of *K*_Monod_ (e.g. the level of *s*_sugar_ for which growth rate is half of its maximum) is not determined solely by the kinetics of the transporter.

To better understand how changes in the model parameters affect the growth rate, we considered a toy example, where all constants are set to a default value of 1 (except for *K*_M;m_ = 2). In Figure 6, we plot the value of *μ* as a function of *s*_sugar_ (based on Eq. (37-38)), each time changing one of the parameters. In almost all cases, we find a trade-off between growth and binding affinity, namely that improving the kinetics (or relaxing a constraint) improves the maximal growth rate, while increasing *K*_Monod_ (i.e. making it worse since higher *K*_Monod_ means higher a concentration of sugar is required to reach the same growth rate). The three exceptions are the response to changes in *ε*_tot_ which only affects *μ*^max^, in *K*_t_, which only affects the *K*_Monod_, and in 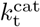, which can improve both growth parameters at the same time. Indeed, by observing the formulae in Eq. (38), one can see that all parameters that are only in the denominator should affect *μ*^max^ and *K*_Monod_ in the same fashion (which leads to a trade-off), while the three parameters in the numerators have each their own unique effect.

**Figure 6.**
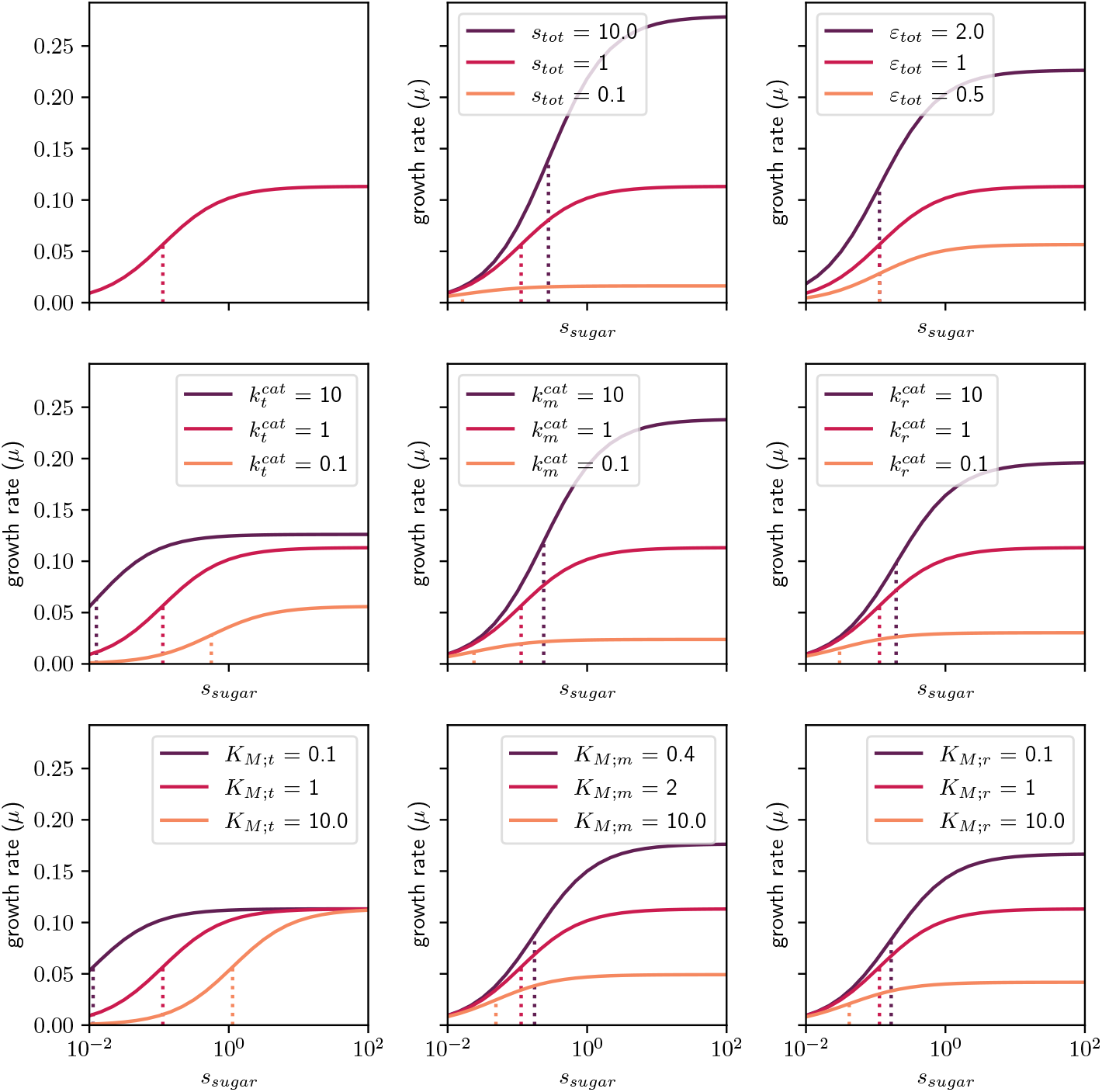
The optimal achievable flux – i.e. the growth rate (*μ*) – for a given total enzyme *ε*_tot_. The model contains three irreversible steps based on Michaelis-Menten kinetics. The default parameters are 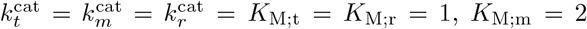 and the upper bounds are set to *s*_tot_ = 1, *ε*_tot_ = 1. The panel on the upper-left corner plots the maximal flux as a function of *s*_sugar_ for the default parameters. In each other panel, only one of the parameters is changed. The vertical dotted lines mark the *K*_Monod_ (i.e. the concentration of *s*_sugar_ where the flux is 50% of the maximum). Note that the x-axis is plotted in log-scale, and therefore the curves have a sigmoid-shape (rather than the familiar Monod-style). See Figure S1 for plots of the enzyme demands.

This result might have implications for the design of optimal enzyme regulation. As the rate of sugar transport is not affected by any of the metabolite concentrations inside the cytoplasm, the transport reaction acts as an information buffer in the sense that any changes in the external sugar concentration do not affect the optimal allocation of enzymes (and metabolites) within the cytoplasm. The only thing that changes is the relative abundance of the transporter versus all other (cytoplasmic) enzymes. This rule becomes clear when plotting the optimal enzyme allocations as a function of the pathway flux (i.e. the growth rate), as shown in Figure 7. The two cytoplasmic fractions (metabolic enzymes and ribosomes) change proportionally and appear as straight lines that cross the axis origin. Since *ε*_tot_ is a constant, the remainder (i.e. the level of the transporter) also decreases linearly with growth rate. These three straight lines extend only up to a point (where *μ* = *μ*^max^). This limit represents the fully saturated transporter, which has an efficiency of 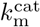 and therefore an optimal abundance of 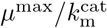. Higher concentrations of *s*_sugar_ would not generate any further benefits.

**Figure 7.**
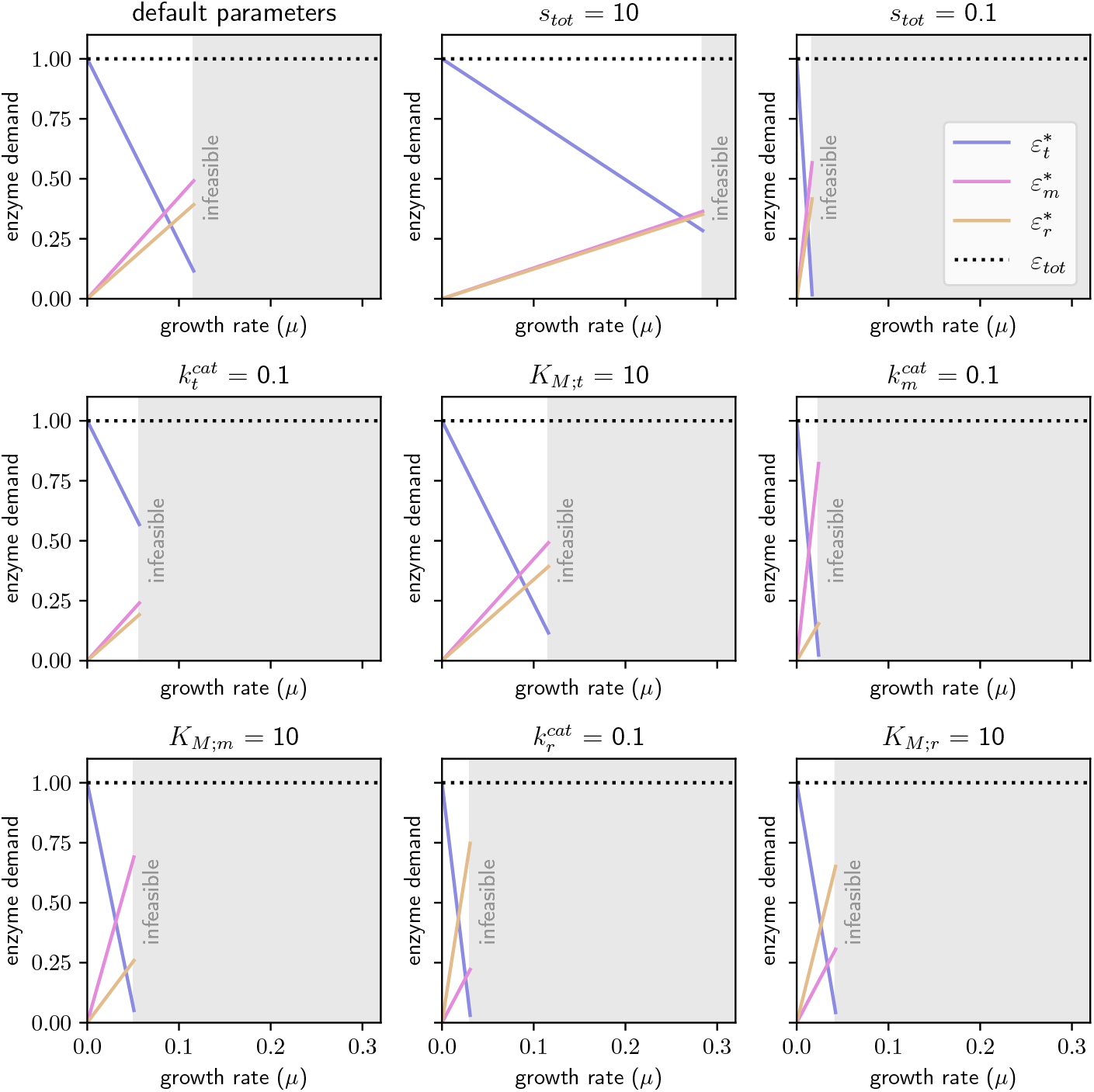
The abundances of the three enzyme sectors (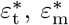, and 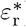) as a function of the flux (*J* ^*∗*^) when the allocation is optimal. The default parameters are 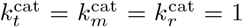, *K*_M:t_ = *K*_M:r_ = 1 *K*_M:m_ = 2 (we chose another value in order prevent the curves for *ε*_m_ and *ε*_r_ from overlapping), and the upper bounds are set to *s*_tot_ = 1, *ε*_tot_ = 1. In each panel only one of these parameters is changed. The gray shaded area marks growth rates that are higher than the maximum, *μ*^max^. The linear protein allocation curves are caused by the irreversible transport step. In the optimization, this leads to constant optimal metabolite concentrations, and the protein fractions 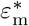 and 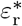 must change proportionally with the flux. The line slopes are given by the enzyme efficiencies at the optimal metabolite concentrations, and the transporter fraction – the residual of the protein budget – must be a falling straight line.

In fact, the principle described above can be shown to be true in a much more general case. As long as one section of the metabolic network is “isolated” from the rest (i.e., connected only by irreversible steps), changes in the upstream parameters will affect the incoming flux, but would not change the optimal allocation of enzymes (and also not the metabolite concentrations). Curiously, in such cases our result predict, based on an underlying kinetic model, that the optimal protein efficiencies are independent of the external glucose concentration and, as a consequence, protein fractions vary linearly. Thus, our model predictions are very similar to ones from previous proteome allocation models that are based on empirical observations.

A series of irreversible steps is only one option for a simple model of cell growth. Alternatively, one might consider cells that live close to chemical equilibrium, in environments that provide only a very small overall thermodynamic driving force. In this case, applying our thermodynamic rate law with the approximation from Eq. (86) yields another simple “growth formula”:

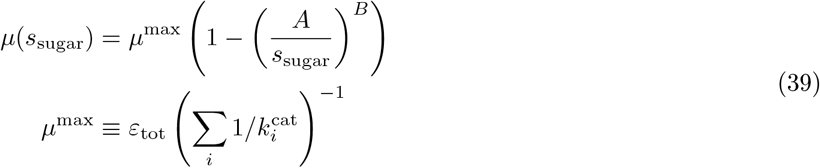

where 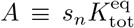 and 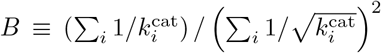. Again, the parameters of this curve depend on all the 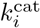 svalues. In contrast to the Michaelis-Menten based growth formula in Equation (38), this formula is also sensitive to product concentrations. This can be interesting because close to chemical equilibrium, product accumulation can decrease growth in a way that cannot be bypassed by any regulatory mechanisms. Instead, cells would profit from “cleaner” strains that remove the accumulating product, which has been proposed as a general mechanism that encourages symbiosis [29].

## 3 Discussion

In this study we addressed the enzyme allocation problem analytically by considering several different rate-law approximations. Along the way, we learned that each approximation comes with its own idiosyncrasies. For instance, a model with Michaelis-Menten kinetics requires an extra metabolite constraint to obtain a valid optimal state, and the *thermodynamic* rate law requires inverting a function in order to find the value of the Lagrange multiplier. Fortunately, there is a very good approximate solution that works well across the entire range of driving forces.

For historical reasons, we solved the optimality problem under the assumption that there is a bound on the sum of enzyme abundances – Σ_*i*_ *ε*_*i*_*≤ε*_tot_ – without explicitly specifying the units. The common interpretation would be that *ε*_*i*_ are *molar* concentrations (where the symbol 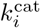 srepresents the turnover numbers in units of 1 over time). However, the total *mass* concentration of all pathway enzymes may be a better proxy for cost since crowding effects in the cytoplasm are often limiting the total mass concentration of all soluble proteins, including most enzymes. Furthermore, fast growing cells are often limited by the rate of protein elongation, and the molecular mass is nearly proportional to the gene length. Therefore, we usually think of *ε*_*i*_ as *mass* concentrations (e.g. in g ×m^*−*3^) and of 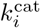 as specific activities (e.g. in mol × s^*−*1^ ×g^*−*1^). Nevertheless, the analytic derivation and solutions provided here are agnostic to the choice of units and are equally valid for both these interpretations. Moreover, one could imagine a completely different set of linear weights for the enzyme cost function (i.e. Σ_*i*_ *w*_*i*_ *ε*_*i*_*≤ε*_tot_, as in Figure 5). If one things of the weights as scaling factors for the *k*^cat^ values, the provided solutions will still hold (while making use of the new “effective” *k*^cat^ values).

Previous studies focused on the “mass-action” rate law, justifying it by saying that the general reversible form derived by Haldane can be approximated at the limit of low concentrations. However, it is quite rare to have all the substrates and products of an enzyme at concentrations that are way below their *K*_M_ values [30]. Furthermore, the limit of all reactants concentrations going to zero is not very meaningful because, in the first place, Haldane derived his rate law assuming enzymes are much less abundant than metabolites [31]. Interestingly, just being close-to-equilibrium is not enough for this approximation, since the product can still affect the rate non-linearly via *η*^kin^ (see Figure 2). Here we tried to consider more comprehensively all the different approximations that yield an analytic solution to optimal enzyme allocation in unbranched pathways (see Table 1).

One of these approximations is the Michaelis-Menten rate law, which is a widely used in enzymatic assays and metabolic modeling of irreversible reactions. Curiously, using it for the simple optimality problem with linear chains leads to paradoxical results: the metabolite concentrations go to infinity, the elasticities vanish, and flux control coefficients are not defined. For solving this problem, we introduced an new upper bound on the total metabolite concentration in order to obtain realistic results and derive analytic solutions based on this rate law. Although it is very reasonable to assume that concentrations of small molecules (and not just enzymes) are restricted in cells, this fact is often ignored in metabolic models. One of the rare cases where this constraint was taken into account is the work of Dourado et al. [32], who found empirical evidence to the fact that there is a balance between enzymes and substrates when minimizing the total mass concentration. In addition to an analytic solution, we also found a new enzyme-control rule for models with a constraint on enzyme and metabolite concentrations: in this case, the enzyme amounts do not only reflect the flux control coefficients, but a sum of flux and concentration control coefficients see Appendix C.2).

Besides “mass-action” and Michaelis-Menten kinetics, we discussed a solution for one other rate law which we call *thermodynamic*, as the rate is only affected by metabolite levels through the thermodynamic driving force (i.e. ignoring any saturation effects). One advantage of this rate law is that it does not require knowing the *K*_M_ values (which are often difficult to come by). The thermodynamic-only approximation was also used for deriving the Max-min Driving Force (MDF) method F.3.5, which similarly aimed to quantify the efficiency of metabolic pathways [33]. But, unlike the solution presented here, MDF does not explicitly optimize a simplified version of a kinetic rate law, but rather applies a heuristic based on the assumption that the lowest driving forces are mainly responsible for increased enzyme demands, and the total driving force should therefore be as evenly distributed as possible. On the other hand, the advantage of the MDF method is that it takes metabolite concentration bounds into consideration.

All throughout this paper, we only considered pathways with uni-uni reactions (one substrate, one product, with stoichiometric coefficients of 1). In reality, many reactions involve cofactor pairs or other substrates or products. Instead of deriving results for this general case, we assumed that these extra reactants exist, but with known concentrations. In this case, the rate laws contain extra terms, but these terms can be rearranged to yield the same simple formulae as in the uni-uni case, but with effective kinetic constants. We demonstrate this for the case of two substrates and two products following convenience kinetics [34] in Appendix F.6. Notably, this logic also applies to reactions with more than two substrates and products as well as to enzyme activation or inhibition with constant activator or inhibitor levels.

This paper can be seen as an exercise in solving the enzyme allocation problem and describing the optimal states analytically using MCA. Although some might argue that the required approximations are not realistic, they do represent a step forward compared to the very common approach of assuming that metabolites have no effect on enzyme efficiency at all. On the other hand, adding metabolite concentrations as extra variables greatly increases the complexity of models and typically renders them unsolvable. Therefore, the solutions provided here might be handy, as the assumption of metabolite steady-state combined with the optimality argument give us analytic expressions that only depend on the initial and final metabolites in the pathway. We demonstrate this result using a toy example for a course-grained model of cell growth, and show how the analytic solutions provide valuable insights about the effects of changes in each parameter – all this without the need to simulate the metabolic network or use non-linear solvers. One could imagine a scenario where metabolic engineers, while trying to improve a biochemical pathway, have to decide in which enzyme to invest their time. Although MCA can perfectly serve this purpose while covering more scenarios, it requires tools and language that can seem opaque to people lacking the proper mathematical background. The solutions provided here should be more approachable and serve as simple guidelines while capturing a wider range of cases than previously explored in the literature. We hope that future studies will apply and extend this approach to other, more complex models.

## Abbreviations

ECM: Enzyme Cost Minimization
PSA: Pathway Specific Activity
MDF: Max-min Driving Force method

## Acknowledgments

This article has originally been conceived as a chapter for the free textbook “Economic Principles in Cell Biology”. A shorter chapter version will be published there. We thank Daan de Groot and Avi Flamholz for their insightful comments on our manuscript.

## A Code availability

All the data, scripts, and resulting plots used in this work are freely available at https://gitlab.com/elad.noor/optimal_enzyme_investment under an MIT open-source license.

## B Symbols

Mathematical symbols are listed in Table S1.

**Table S1:** The names of the variables and constants used in this work, along with their mathematical symbol.

| Parameter name | symbol |
| --- | --- |
| Turnover constant | $k^{\text{cat}}$ |
| Turnover constant (forward) | $k_+^{\text{cat}}$ |
| Turnover constant (backward) | $k_-^{\text{cat}}$ |
| Thermodynamic equilibrium constant | $K^{\text{eq}}$ |
| Michaelis-Menten constant | $K_M$ |
| Michaelis-Menten constant (substrate) | $K_S$ |
| Michaelis-Menten constant (product) | $K_P$ |
| Thermodynamic Driving Force | $\theta$ |
| Reaction rate | $v$ |
| Pathway flux | $J$ |
| Enzyme concentration/demand | $\varepsilon$ |

## C Revisiting the enzyme-control rule

### C.1 Why the enzyme-control rule fails with Michaelis-Menten kinetics

The enzyme-control rule fails in a very simple case, the unbranched metabolic pathway with Michaelis-Menten rate laws. We can easily see this. Since the rate law is irreversible, the rate in each reaction depends only on the substrate and enzyme level (and not on anything that happens downstream). In steady state, all reaction rates are fixed by the stationary flux. Taken together, this means that the flux depends only on the initial pathway substrate and the level of the first enzyme, so this enzyme has full flux control and all other enzymes have zero control. However, according to the enzyme-control rule, this would mean that the first enzyme is the only enzyme that should be expressed. This is obviously wrong, because without the other enzymes, no steady flux would be possible. So how can this be?

In fact, optimal steady states in models with irreversible rate laws have some pathological properties. To maximize enzyme efficiency, all metabolite levels must go to infinity, the enzymes are completely saturated, and the metabolite elasticities are all zero. As a result, the flux in every reaction is given by 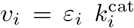, so the stationary flux completely determines all enzyme levels. If an enzyme level deviates from its prescribed value, its reaction rate will no longer match the other rates and the steady state breaks down^2^. Now we can see what went wrong: in the optimal state, any small change of an enzyme level will break the steady state; this means that the control coefficients are not even defined, and the enzyme-control rule cannot apply. All this does not only occur with the Michaelis-Menten rate law, but with any irreversible, saturable rate law.

In the case of the Michaelis-Mention rate law, the practical solution to this problem is simple. Since the optimal state requires infinite metabolite concentrations, and since this is not realistic, we need to change our model. We may either consider slightly reversible rate laws (which would penalize high product concentrations) or we introduce an extra density constraint that penalizes high metabolite levels.

### C.2 An enzyme-control rule for models with general density constraints

One way to obtain meaningful optimal states, even with Michaelis-Menten kinetics, is to constrain the metabolite concentrations. The new optimality problem leads to different solutions and to a different enzyme-control rule.

Aside from high enzyme levels, also high metabolite concentrations may put a burden on cells, and considering this burden will lead to different solutions for enzyme levels. Setting up optimality problems with such a constraint raises a couple of questions: should we use separate constraints on enzyme levels and on metabolite concentrations, or just one density constraint on their sum? How should the different compounds be weighted? In fact, the resulting optimality problems are not very different, and we demonstrate here one specific approach: we describe all compounds (enzymes and metablites) by their concentrations and consider a bound on their weighted sum (where weights may have different meanings, for example molecular masses if the aim is to restrict the total mass density in the cell).

We thus consider the optimality problem

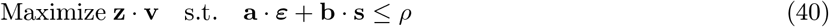

where **v, s**, and ***ε*** denote reaction fluxes, metabolite concentrations, and enzyme levels, and where stationarity (of **v**) and rate laws (between **v, s**, and ***ε*** must be satisfied as constraints. The maximization objective is a linear benefit function **z ·v** scoring the fluxes with linear weights *z*_*i*_, while the constrained density is linear in all the concentrations, with linear weights^3^ *a*_*l*_ (for enzymes) and *b*_*i*_ (for metabolites), e.g. representing effective molecule sizes^4^. To derive the enzyme-control rule, we now reformulate the optimality problem with enzyme levels as the only free variables, and steady-state fluxes and concentrations depending on them (satisfying stationary and the rate laws). We obtain

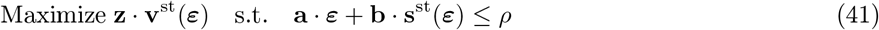

This optimality problem leads to a new enzyme-control rule (proof in Appendix F.7)

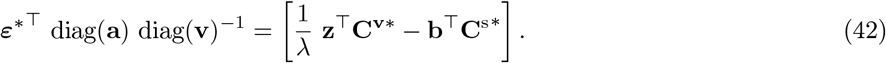

In the general case (network with non-uniform flux distribution **v**, a more complicated flux benefit function with derivatives 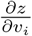, and individual weights *a*_*l*_ and *b*_*i*_ for all enzymes and metabolites, all these vectors would enter the formula as weights.

In a simple linear pathway, our rule can be simplified. We assume the same steady-state flux *J* in all the reactions, a benefit function given by *J*, and a density constraint with equal weights *a* for all enzymes, and equal weight *b* for all the metabolites. In this case, Eq. (42) simplifies to

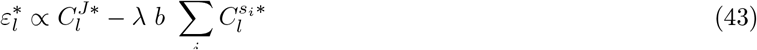

which replaces the enzyme control rule (8). Given the sum of enyzme levels in the optimal state, called 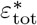, we can also write this as (see F.7)

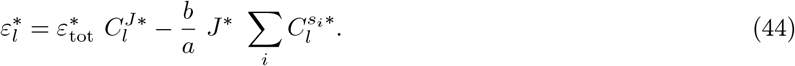

Since the Lagrange multiplier *λ* arises from an upper bound in a maximization problem, it is negative in the optimal state, and the second term including the minus sign will be positive.

Compared to the enzyme-control rule (8), the right hand side now contains an extra term: it is given by the metabolite control coefficients, summed over all metabolites, and with a known prefactor. If the ratio of cost weights *b/a* is small compared to the flux, this term can be neglected and we obtain again our original enzyme-control rule.

### C.3 A sufficient condition for stable states in unbranched pathways

Another possible reason why control coefficients may not be defined are unstable metabolic states. A metabolic steady state can be dynamically stable or unstable. In an unstable state, even small perturbations or chemical noise would drive the system far away from its original state. In MCA we are interested in stable states, because unstable states are not able to persist even under small chemical noise in the cell, and because mathematically, control coefficients for unstable states are not even defined. A metabolic state is asymptotically stable if the Jacobian **N E**_c_ in this state has only negative eigenvalues. But what does this mean in practice? In Appendix F.8, we derive a sufficient (but not necessary) condition for stable steady states in unbranched metabolic pathways, based on Gershgorin’s theorem: A steady state in an unbranched metabolic pathway is stable if (but not only if) the first reaction 1 is not completely irreversible (i.e. it has a non-zero product elasticity), the last reaction is not completely saturated (i.e. it has a non-zero substrate elasticity), and in all reactions in between, the substrate elasticity is larger than the (absolute) product elasticity. This condition is not just a chek for our models, but it also describes a potential way in which cells could achieve stability. For a reversible mass-action rate law, the latter condition is satisfied if 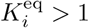, and the first two conditions are generally satisfied.

## D The enzyme-elasticity rule

The connectivity theorem for flux control coefficients

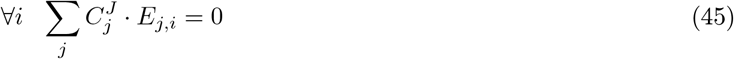

relates the flux control coefficients 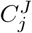 (between enzyme *ε*_*j*_ and the pathway flux *J*) to the unscaled elasticities 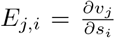 (between metabolite *s*_*i*_ and reaction rate *v*_*j*_). In optimal state, the enzyme-control rule tells us that control coefficients and enzyme levels must be proportional. Therefore we can replace 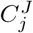 with *ε*_*j*_ in the equation above. For each metabolite *i*, we obtain an equation

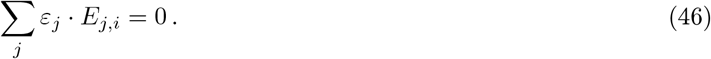

In a linear pathway, each metabolite has only one producing and one consuming reaction (with indices *i* and *i* + 1, and we obtain the equality

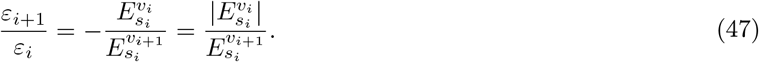

In the last step, we used the fact that the backward (product) elasticity 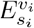 is negative and the forward (substrate) elasticity 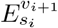 is positive.

**Example: Mass-action rate law** With the mass-action kinetic rate law in the form *v*_*j*_ = *ε*_*j*_ *·* (*k*_*j*_*s*_*j−*1_ −*k*_*−j*_*s*_*j*_) (see Appendix F.4), Eq. (48) becomes:

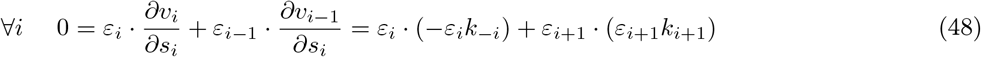

which by rearranging leads us to the relationship:

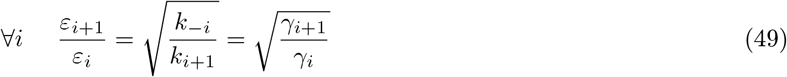

(where we remind ourselves of the definition 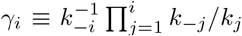. Applying this equation recursively, we can see that 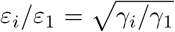, which means that *ε*_*i*_ is proportional to 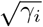. Since we know that the sum of all enzyme demands gives *ε*_tot_, we can obtain the explicit formula for *ε*_*i*_:

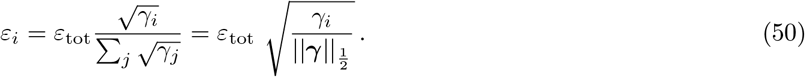

This is the exact same result we got by directly optimizing the pathway flux (see Equation (108) in Appendix F.4).

**Figure S1:**
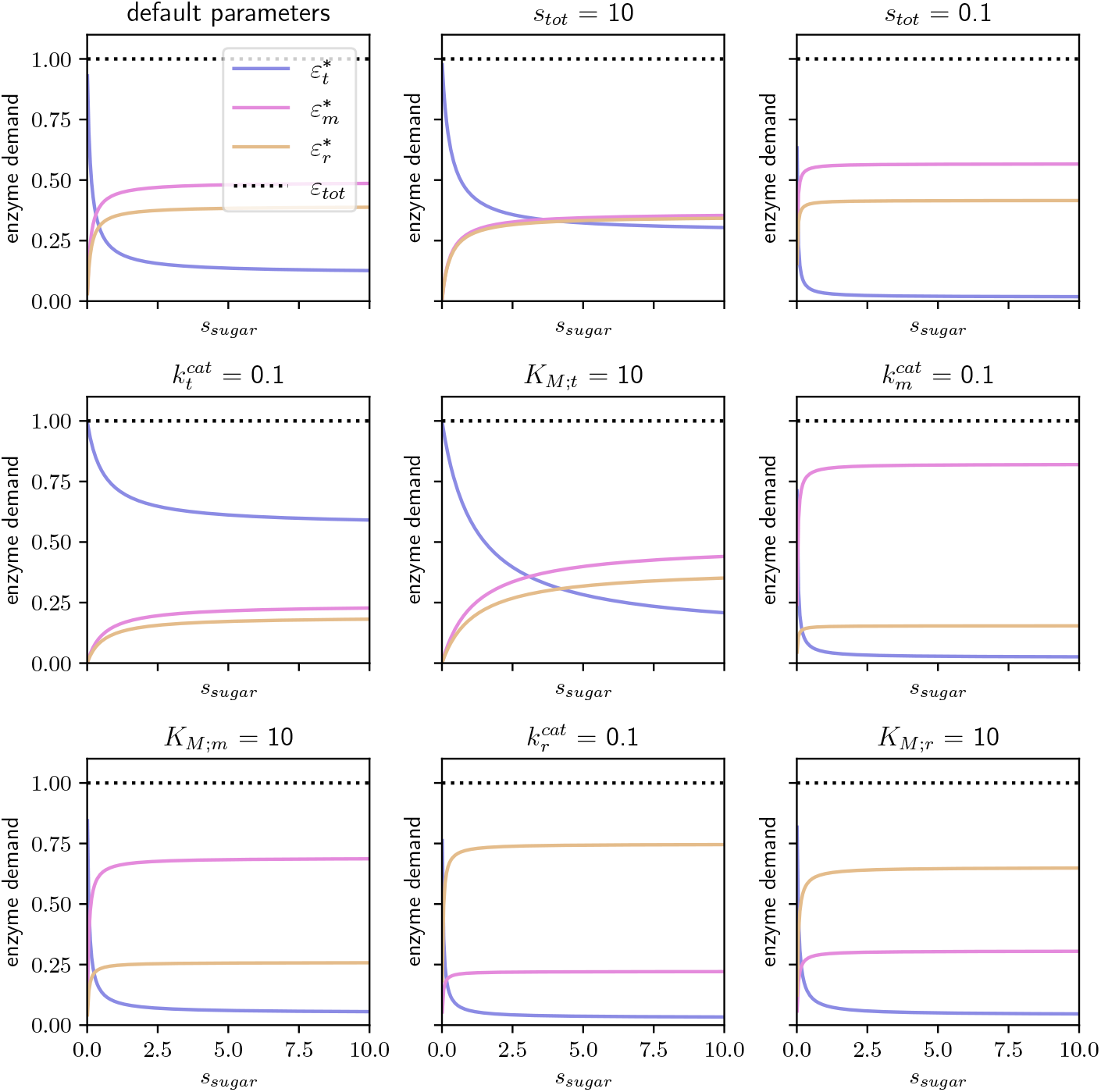
Enzyme demands in a simple cell model, depending on external sugar concentration. The abundances of the three enzyme sectors, when the allocation is optimal, as a function of the external sugar concentration (*s*_sugar_). The default parameters are 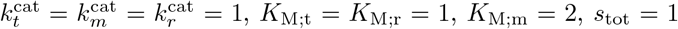, and *ε*_tot_ = 1. In each panel only one of these parameters is changed.

## E A cell model with enzyme kinetics and metabolite constraints

The cell model is described by three irreversible reactions:

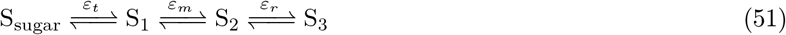

The optimal achievable flux (*J* ^*∗*^) for a given total enzyme *ε*_tot_ is given in Eqs (37-38). Figure S1 shows the individual enzyme demands for the three steps, as a function of the external sugar concentration and *J* ^*∗*^:

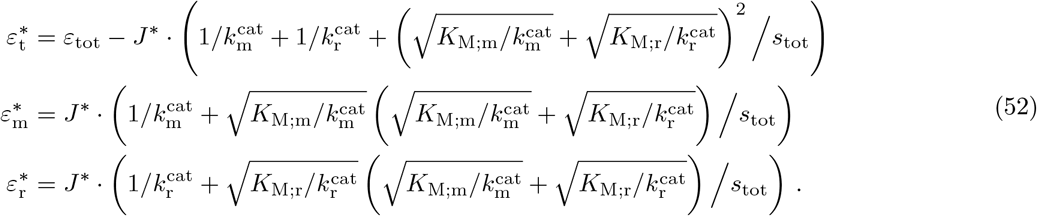

Note that also 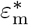 and 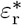 change with *s*_sugar_ even though it does not explicitly appear in the equations, because *J* ^*∗*^ itself is a function of *s*_sugar_.

## F Mathematical derivations and proofs

This appendix contains many of the detailed mathematical proofs and derivations used in the manuscript for metabolic steady-states, minimum enzyme demand / maximal flux, connectivity theorem, and enzyme control rules. The derivations are grouped by the corresponding rate law in the following order: (1) Michaelis-Menten; (2) Thermodynamic; (3) Reversible mass-action; (4) Haldane. An overview of useful equations is given in Table S2.

**Table S2:** Overview of formulae for steady-state variables and control coefficients. This table summarizes formulae for unbranched metabolic pathways with different rate laws (rows). The columns refer to steady-state concentrations and flux (as a function of enzyme levels and metabolite concentrations); steady-state concentrations as a function of enzyme levels, pathway substrate concentration, and steady-state flux (but not of the product concentration); reaction elasticities; and concentration and flux control coefficients.

| | $s_i^{\text{st}}(s_0, s_n, \epsilon)$ | $J(s_0, s_n, \epsilon)$ | $s_i(s_0, J, \epsilon)$ | Elasticities | $C^s$ | $C^J$ |
| --- | --- | --- | --- | --- | --- | --- |
| Michaelis-Menten | Eq. (55) | $v_1(s_0)$ | Eq. (56) | Eq. (54) | | Eq. (57) |
| Thermodynamic (const $\eta^{\text{kin}}$) | Eq. (34) | | Eq. (34) | | Eq. (94) | |
| Thermodynamic (substrate sat) |  |  |  |  |  | sec F.3.2 |
| Mass-action | Eq. (26) | Eq. (103) |  | Eq. (111) |  | sec F.4.2 |
| Haldane | - | - | Eq. (19) in [17] | e.g. [35] | - | - |

### F.1 Michaelis-Menten kinetics

The Michaelis-Menten rate law is an approximation of the Haldane rate law for irreversible reactions:

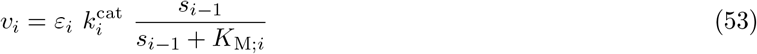

with reactant elasticities

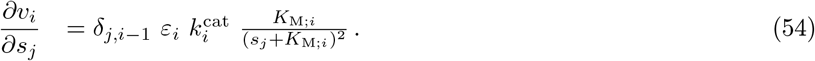

#### F.1.1 Steady-state concentrations

Computing the steady-state metabolite concentrations in a pathway with Michaelis-Menten kinetics is possible by inverting Eq. (53).

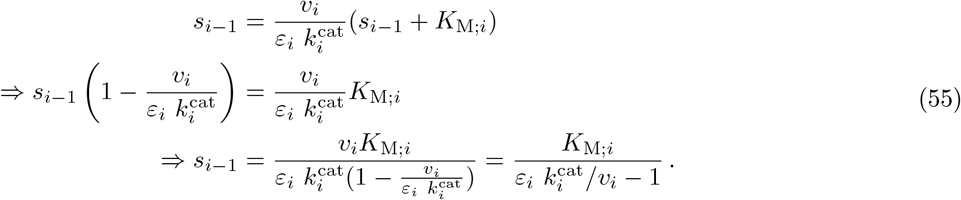

The steady-state concentrations as a function of the pathway flux can be found by replacing *v*_*i*_ with *J* for all reactions. For any two reactions (*i* and *j*), we can relate between their substrate levels by replacing *v*_*i*_ in Eq. (55) with *J* and using the rate law for *J* = *v*_*j*_ from Eq. (53)

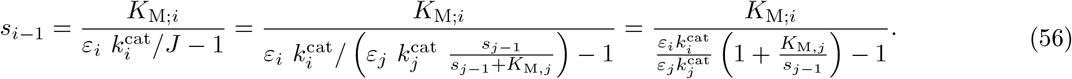

In fact, applying this condition to all metabolites is a sufficient condition for having a steady-state. Specifically, if we start with a given concentration *s*_0_ and an enzyme abundance vector ***ε***, there is only one solution for the steady-state flux and for the metabolite levels (**s**) which support it.

#### F.1.2 Flux control coefficients

We can compute the flux control coefficients directly by taking derivatives: given that *J* = *v*_1_ and that *v*_1_ depends only on the external substrate, described by a fixed parameter and therefore not dependent on the downstream *ε*_*i*_, the flux control coefficients read

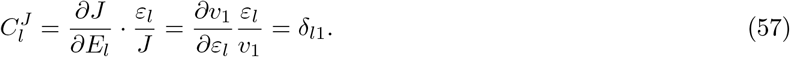

As expected, due to the irreversible rate law, only the first enzyme can have control (except for the pathological case where metabolites concentrations become infinite and flux control is not defined).

#### F.1.3 Minimum enzyme demand / maximal flux with a metabolite constraint

At steady state (with flux *J*), Eq. (53) can also be used to express the enzyme demands:

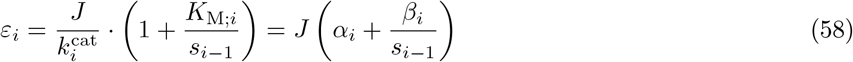

where we defined 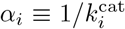 and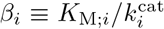 as before. Now we can equate the sum of all enzyme demands to *ε*_tot_:

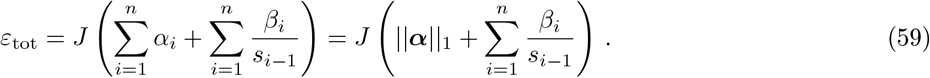

This inverse relationship between *ε*_tot_ and the metabolite concentrations means that in order to minimize *ε*_tot_ for a given flux, all *s*_*i*_ should be as large as possible. Obviously, infinite concentrations are not feasible. To avoid non-physical solutions, we can add a constraint on the total concentration of metabolites 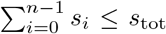. Now, using Lagrange’s method, we define 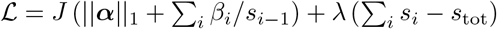 and take the derivatives:

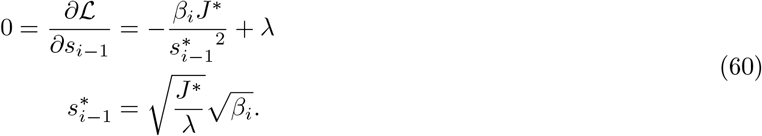

To find *λ*, we can now use the constraint on the sum of all metabolites (assuming the upper bound is hit):

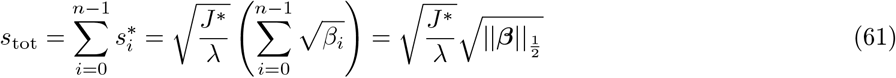

where ***β*** is the vector of all *β*_*i*_, and 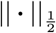 is the l_1*/*2_ norm. Therefore, we can replace 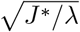 with 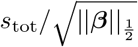 in Eq. (60) and find an explicit expression for 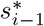:

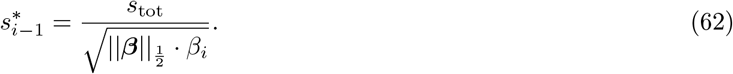

By combining the results from Eqs (59) and (62), we can see that:

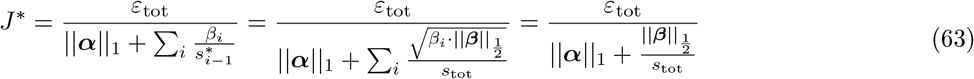

and similarly we can also find the optimal enzyme levels:

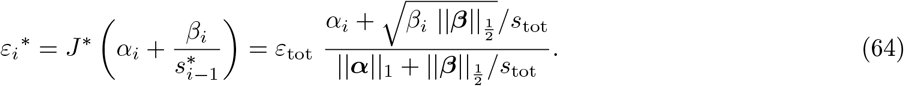

#### F.1.4 Case where the first substrate has fixed concentration

If our pathway model is used to represent a cell growing on an external substrate (S_0_), then it would be unrealistic to assume that the concentration *s*_0_ is subject to optimization. Instead, we can assume it is fixed (e.g. based on environmental conditions) and that the cell optimizes only the concentrations of enzymes and internal metabolites.

From Eq. (63) we can see that:

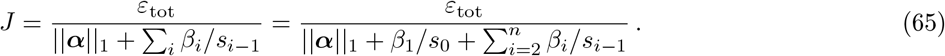

Since *s*_0_ is fixed, we consider the term *β*_1_*/s*_0_ to be part of the constant part of the denominator (usually just 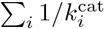. Furthermore, we can redefine 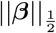 to exclude the value for the first enzyme (*β*_1_) and impose the upper bound *s*_tot_ only on the sum of internal metabolites. In this case, the solution we got in Eqs (60)-(61) will still be applicable, and therefore we can write:

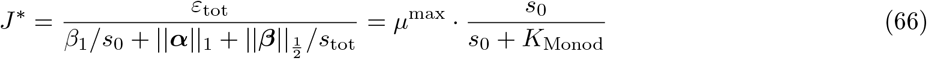

where we defined

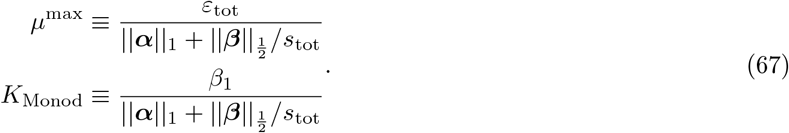

As we can see, the solution provides a prediction for the Monod constant – i.e., the concentration of substrate (*s*_0_) where the growth rate reaches half of its maximum. If the transporter turnover constant 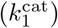 is indeed much slower than in all other enzymes,its control coefficient is 1 and we will get *K*_Monod_ = *K*_M,1_, which is often naïvely assumed. However, if the control is distributed along the pathway, the Monod constant can become significantly smaller than *K*_M,1_.

We also obtain expressions for the individual enzyme levels, using Eq. (64). Because *s*_0_ is fixed, the demand for the first enzyme is can be simply expressed as:

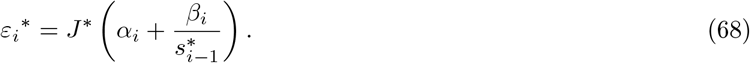

For the first enzyme, the substrate (*s*_0_) is fixed so the expression yields simply *ε*_1_^*∗*^ = *J* ^*∗*^ (*α*_1_ + *β*_1_*/s*_0_), while for all other enzymes (*∀i >* 0) we can use the solution for 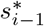 from Eq. (62), i.e.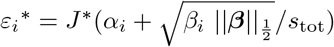.

### F.2 Thermodynamic rate law with fixed kinetic efficiency

The thermodynamic rate law with fixed kinetic efficiency reads:

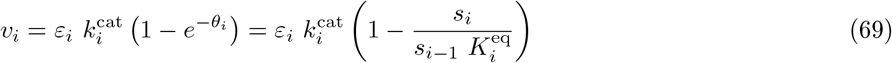

where 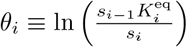

#### F.2.1 Steady-state concentrations

Assuming a steady state (i.e. all fluxes are equal to *J*), we can find a simple recursion formula for the concentrations *s*_*i*_:

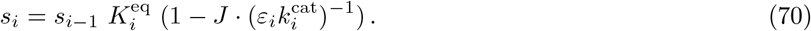

By solving the recursion for *s*_*n*_ we get:

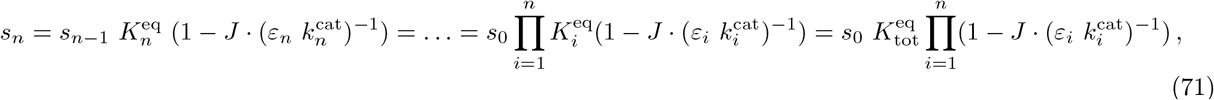

where we used the fact that 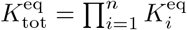.

The driving force of the entire pathway, which we denote by *θ*_tot_, is the sum of all the individual driving forces:

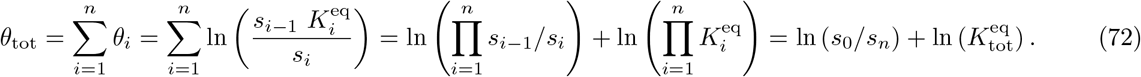

Since *θ*_tot_ depends only on 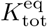, *s*_0_, and *s*_*n*_ – all of which are constant – it is a constant as well. Therefore, we can use this result to rewrite Equation (71) as:

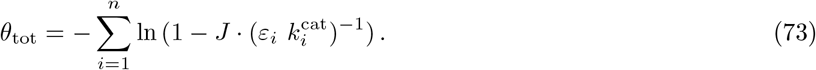

#### F.2.2 Minimum enzyme demand / maximal flux

Directly solving the flux maximization problem using Equation (73) would be difficult, because it cannot be solved analytically to find an expression for *J* as a function of enzyme levels. However, it is rather simple to do the opposite and express the enzyme levels as functions of *J* and the other parameters, sum them up and compare to the total (*ε*_tot_):

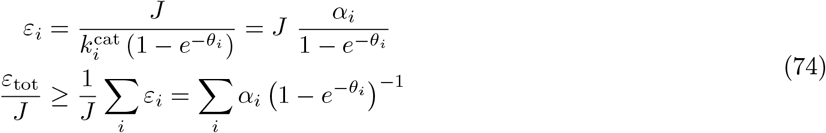

where we define 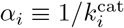 for simplicity.

Minimizing *ε*_tot_*/J* is an optimization problem that can be solved directly using the *s*_*i*_ as variables, but it can be easier to solve if we consider the *θ*_*i*_ to be the independent variables instead. This variable switch is justified because there is a linear homomorphism between the two sets 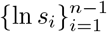 and 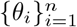, given fixed *s*_0_ and *s*_*n*_ and under the constraint that Σ*θ*_*i*_ = *θ*.

Now, we want to find an optimal set of values for the driving forces, denoted 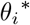, that sum up to *θ* and maximize *J* (similarly, *J* ^*∗*^ and 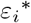 would be the corresponding values of *J* and *ε*_*i*_ at the optimum).

##### Lemma F.1.

*The values of x_i_ that minimize the function* 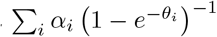 *under the constraint* Σ_*i*_*x*_*i*_ = *x*_*tot*_, *satisfy*

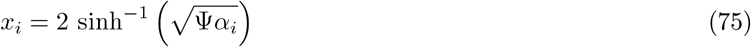

*for some* Ψ ∈ ℝ.

*Proof*. Using Lagrange’s method, we first define 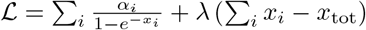

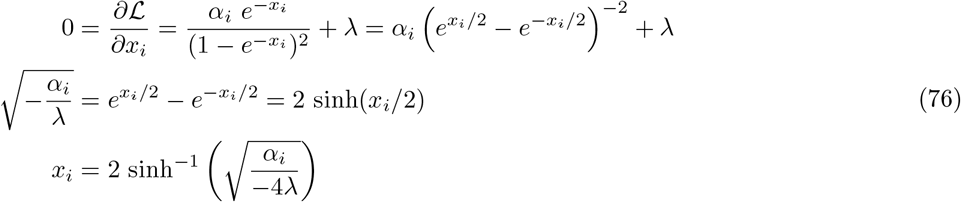

and if we define 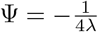, we can see that it proves the lemma.

Using Lemma (F.1), one can see that the optimal distribution of driving forces satisfies the following relationship, for some value of Ψ

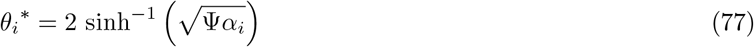

where the exact value of Ψ can be determined by applying the constraint on the sum of *θ*_*i*_ from Eq. (72):

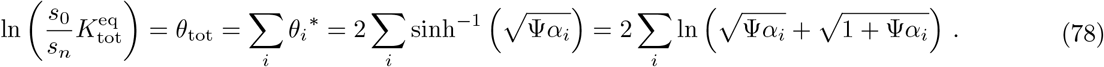

Unfortunately, there is no analytical solution to this equation. Nevertheless, the function on the right-hand side is strictly monotonically increasing with Ψ (in the entire range Ψ ∈ℝ), so solving it numerically should be straightforward. Moreover, we can still express 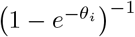 as a function of Ψ (see Appendix F.2.2):

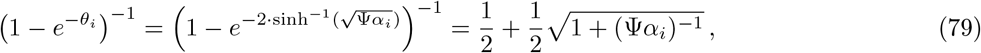

where we use the fact that 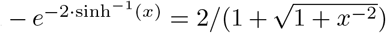. Therefore, the solutions for the maximal flux and individual enzyme allocations would be:

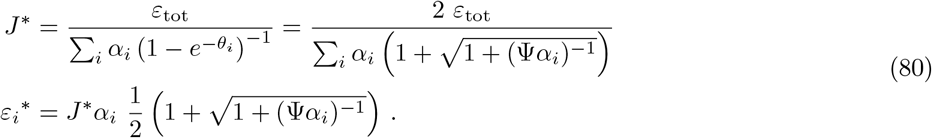

#### F.2.3 Limit of small driving forces

If we further assume that the total driving force *θ*_tot_ is small (and therefore also each one of the reaction driving forces *θ*_*i*_ is even smaller), then we can write:

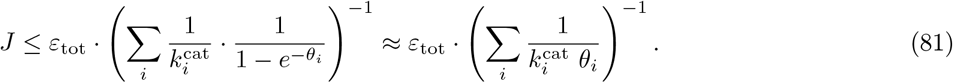

As we did before, the solution for *J* ^*∗*^ can be found by minimizing *ε*_tot_*/J* under the constraint Σ_*i*_ *θ*_*i*_ = *θ*_tot_, using the Lagrange method: 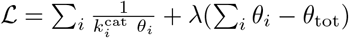.

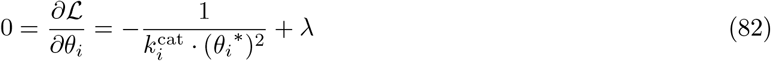

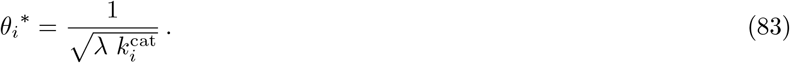

By applying the constraint 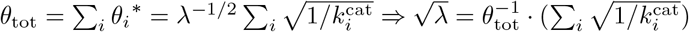, we can write:

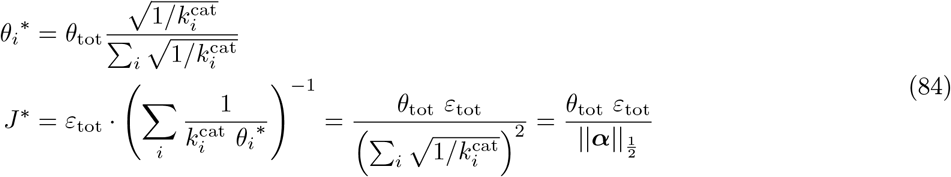

and furthermore, we can see that:

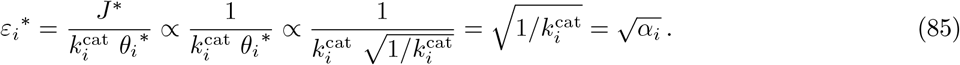

### F.3 Approximate analytical formula

Based on the two limits for values of the total driving force (*θ*_tot_ *→* 0 and *θ*_tot_ *→ ∞*), we can construct an approximation that converges with these limits for extreme values, following the template 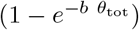.

The solutions for the two limits are:

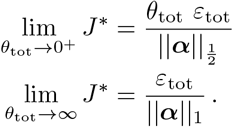

One can see that the approximate formula that has the same limits will be:

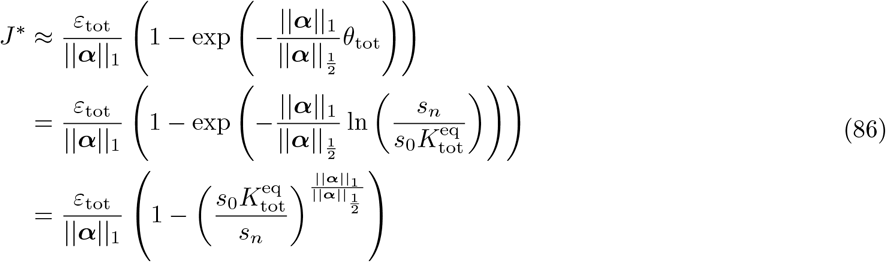

Figure S2 illustrates how this approximation holds up against the precise solution (which is based on the closed-form formula, but requires inverting Eq. (78) numerically to find Ψ). One can appreciate that the approximation is a very good one across all values of *θ*_tot_ and for all choices of *k*^cat^ values tested here. In Figure S3, we compare the approximation to solutions found numerically using convex optimization, i.e. directly optimizing *J* over the space of metabolite concentrations using a convex optimization solver (CVXPY [36, 37]).

**Figure S2:**
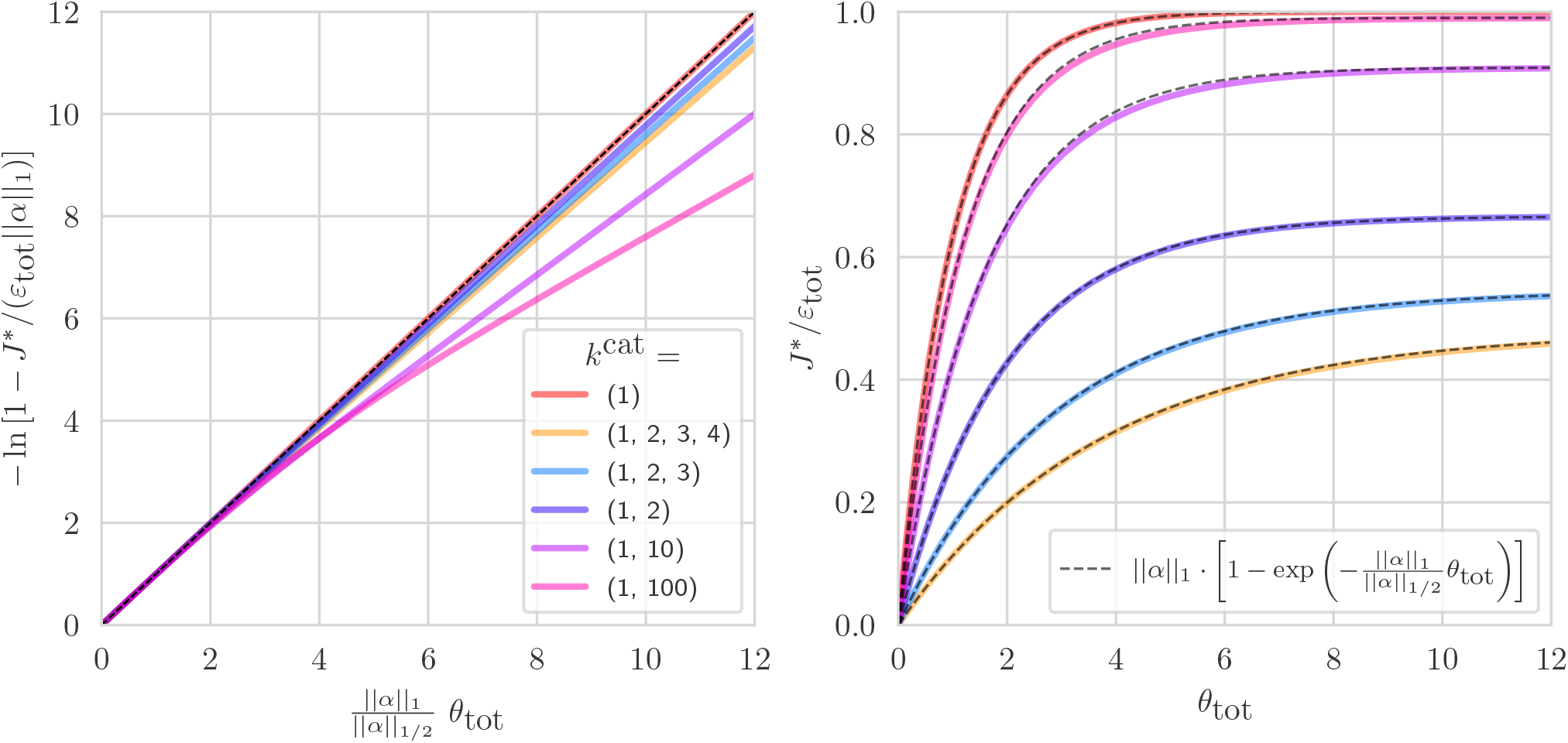
Approximate formula for the thermodynamic rate law case. Left panel: by re-scaling the x-axis by a factor of ||*α*||_1_*/*||*α*||_1*/*2_ and transforming the y-axis using *y* = −ln (1 −*J* ^*∗*^*/ε*_tot_ ·||***α***||_1_), the precise solutions based on Eq. (80) appear almost linear. Each curve represent a different set of *k*^cat^ values (and a varying number of enzymes in the pathway, between 1-4). The approximation based on Eq. (86) is depicted by the dashed black line, which goes exactly along the diagonal for this set of axes. In the right panel, we plot the same precise solutions and approximations as in the left panel, but without transforming the axes.

**Figure S3:**
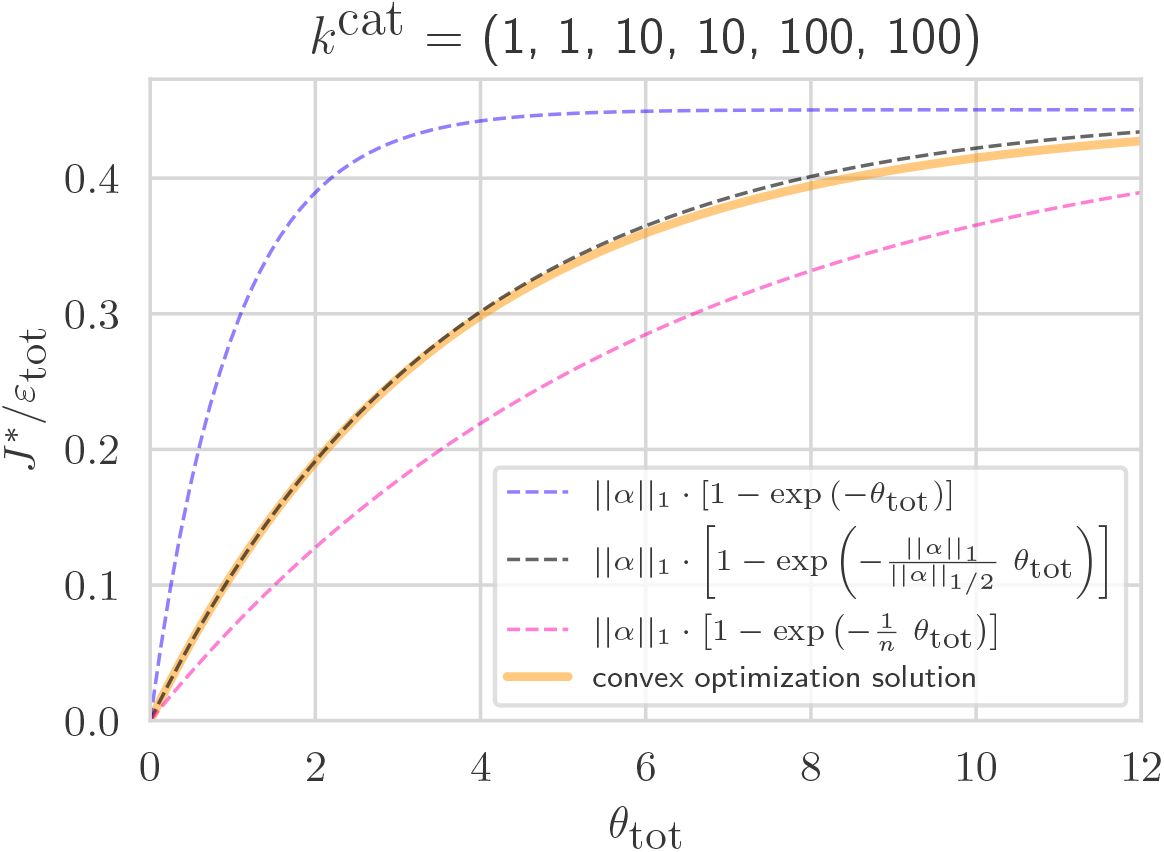
Comparing the approximate solution for the thermodynamic rate law case to the numerical solution based on convex optimization. To arrive at the convex optimization solution (orange curve) we used CVXPY [36, 37]. The precise solution (based on the analytical formula from Eq. (80)) overlaps completely with the one using convex optimization and therefore is not shown. The dashed black line represent the approximate solution. In addition, we plot two other “naïve” approximations: a lower bound (pink dashed line) which represent the rate assuming that the driving force *θ*_tot_ is uniformly distributed along the pathway – i.e. for every reaction *θ*_*i*_ = *θ*_tot_*/n* – and an upper bound (violet dashed line) which represents a case where all the driving force is concentrated in one step, while all other steps have negligible cost.

#### F.3.1 Connectivity theorem

The unscaled metabolite elasticities for the thermodynamic rate law read

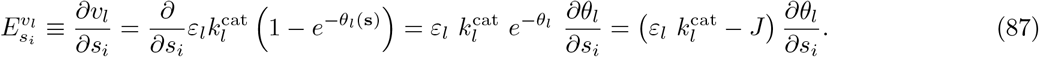

In the last step we used the fact that 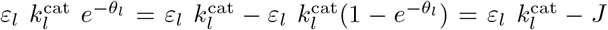. With flux control coefficients proportional to 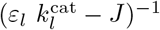, we can now derive the connectivity theorem for each metabolite *s*_*i*_,

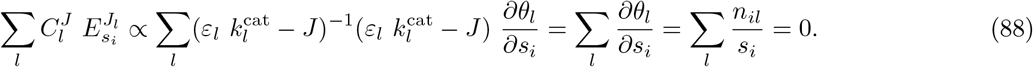

Here we used 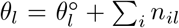 ln *s*_*i*_ with derivative 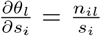 (in the second but last step), and the fact that our reactions are uni-molecular, with stoichiometric coefficients of *±*1 for substrate and products (in the last step).

#### F.3.2 Flux control coefficients

We start by turning Eq. (71) into an equality constraint:

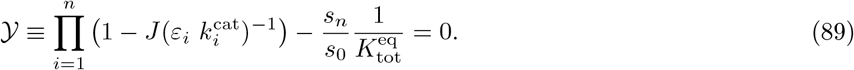

This equation implicitly determines the flux *J* (given the enzyme levels *ε*_*i*_ and the external concentrations *s*_0_ and *s*_*n*_). Since 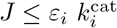, all product terms are positive and decreasing in *J*, and so the entire product is a decreasing function of *J*. Therefore, there can be only one solution with a positive flux. While Eq. (89) cannot be solved for *J* directly, it determines *J* implicitly and is sufficient for computing the flux response coefficients. For doing this we use the following lemma:

##### Lemma F.2.

*We consider two variables x and y, constrained by an equality f* (*x, y*) = 0. *We assume that for each given value of x, the constraint is satisfied by a single value of y, which we call y* = *g*(*x*). *To compute the derivative ∂g*(*x*)*/∂x directly from f* (*x, y*) *without an explicit expression for g*(*x*), *we insert g*(*x*) *and write the constraint as h*(*x*) = *f* (*x, g*(*x*)) = 0. *With the chain rule, we can write this as* 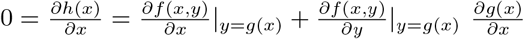, *and obtain*

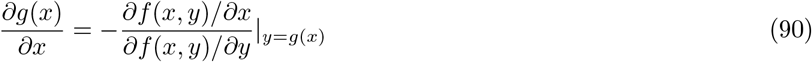

To compute the flux control coefficients, we now see Eq. (89) as a constraint between an enzyme level *ε*_*i*_ and th eflux *J*, assuming that all other enzyme levels are given. This constraint implicitly defines a function *J* (*ε*_*i*_). To compute the derivative *∂J/∂ε*_*i*_ (assuming all other enzyme levels are constant), we set *x → ε*_*i*_; *y → J* ; *g*(*x*) *→ J* (*ε*_*i*_); and *f* (*x, y*) *→ Y*(*ε*_*i*_, *J*), and obtain:

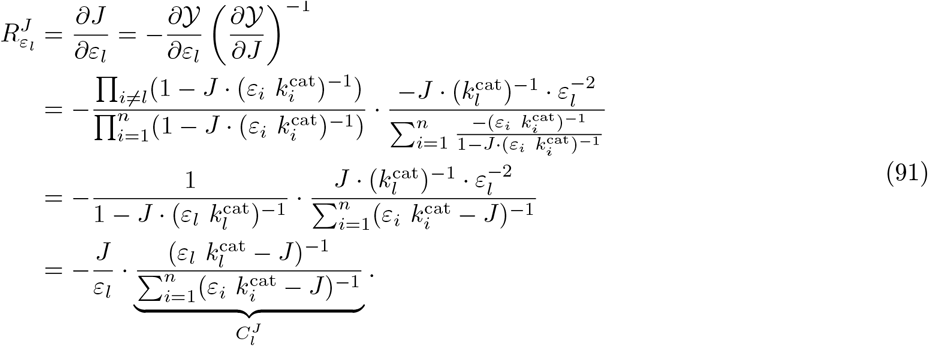

The result is simple: the control coefficients 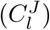 are proportional to 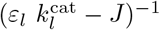 and sum to 1 as required by the summation theorem.

#### F.3.3 Concentration control coefficients

Using Eq. (71), which we can solve for any *s*_*j*_ in the same way as for *s*_*n*_, we obtain 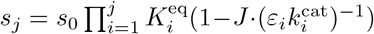 and therefore

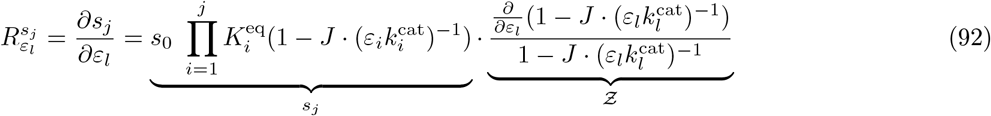

for all enzyme indices *l ≤j* (otherwise 0). The first product term is simply *s*_*j*_. The second product term can be rewritten as:

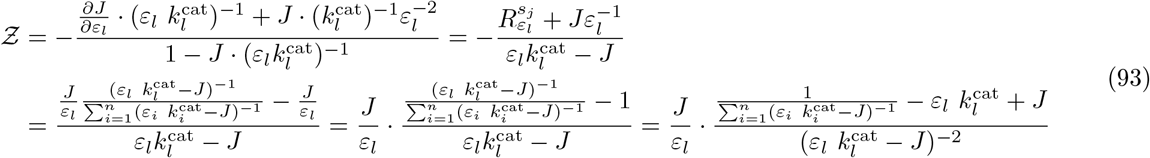

Putting all this together, we obtain the concentration control coefficients

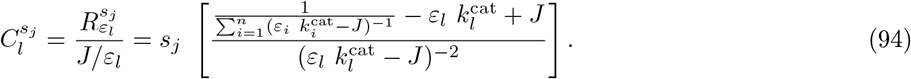

Again, this formula holds only for *l ≤ j* (otherwise 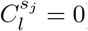).

#### F.3.4 Enzyme-control rule

To show that the control coefficients Eq. (35) satisfy the enzyme-control rule, we assume that the rule is correct, insert our formula for control coefficients, derive from it an expression for the enzyme levels, and compare the result to our formula Eq. (80). We do this now step by step. Starting from the proportionality

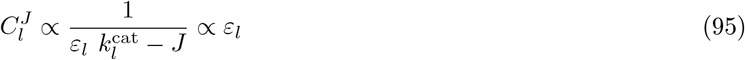

we obtain

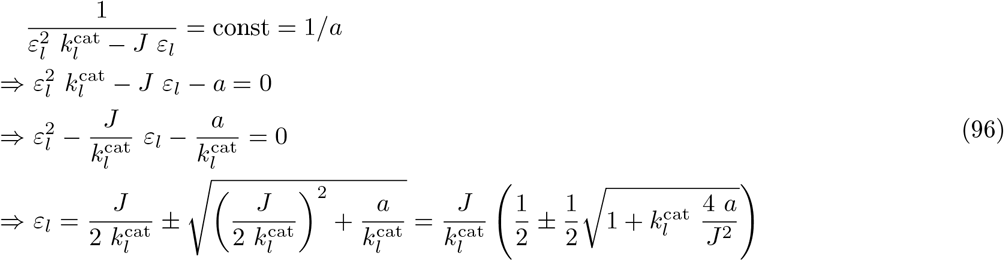

The solution with a minus sign is physically meaningless and can be discarded. Remember that we defined 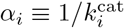, so we can rewrite Eq. (96) as

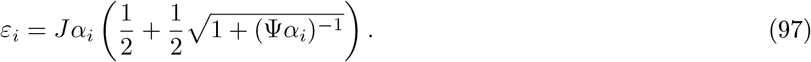

Here we also replaced *J* ^2^*/*(4*a*) with Ψ by comparing this solution to Eq. (80) (and using the fact that the value of this constant is uniquely determined by the constraint Σ_*i*_ *ε*_*i*_ = *ε*_tot_).

#### F.3.5 A note on Max-min Driving Force

The Max-Min Driving Force method (MDF) is a heuristics for finding realistic metabolite concentrations in metabolic models with known flux directions. It relies on the fact that very small driving forces lead to large enzyme demands and should be avoided, while very large driving forces do not further improve the enzyme demand. The MDF heuristics can be seen as a way to reduce enzyme cost. To see this, we start from the thermodynamic rate law and consider a lower bound on the enzyme demand:

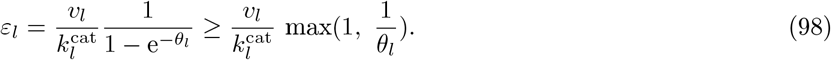

where we use the fact that *e*^*−x*^ *≥* max(0, 1 *− x*). We now consider again an unbranched pathway with a steady forward flux and a given overall driving force *θ*. In this case, we obtain the constraints *θ*_*l*_ *≥* 0 and Σ_*l*_ *θ*_*l*_ = *θ*. The forces *θ*_*l*_ are not free variables, but a function 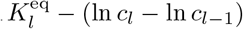 of the metabolite concentrations, which in turn are constrained by physiological ranges. Let us see how the MDF principle can be derived from our present optimality problem. Since pathway flux and total enzyme demand scale proportionally at given metabolite concentrations, a flux maximization at a fixed enzyme budget is equivalent to minimizing the total enzyme demand Σ_*l*_ *ε*_*l*_ at a given flux. With Eq. (98), we can approximate this demand as

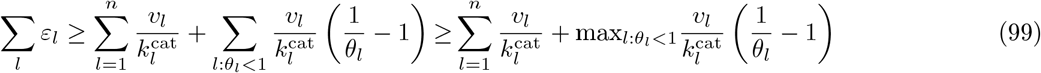

where *l* : *θ*_*l*_ *<* 1 denotes the indices of reactions with driving forces smaller than 1. The first term describes the constant enzyme demand of a hypothetical model in which all enzymes work at their maximal speed. The second term concerns reactions with driving forces smaller than 1 (if such reactions exist) and denotes the maximal value of 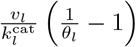 among these reactions. To minimize the overall enzyme demand, we choose the heuristics of minimizing this term. This is equivalent to maximizing

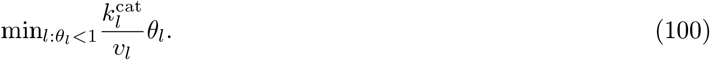

where *l* runs again over all reactions with a driving force smaller than 1. Thus, we postulate that the smallest driving force in the pathway (weighted by 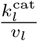) should be as large as possible. The original MDF driving force method employs some additional simplifications. First, we neglect the prefactor 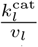 (assuming that nothing is known about enzyme kinetics). Second, if all the reactions have driving forces larger than 1, we let the index *l* run over all reactions, assuming that it will still be the smallest driving force that causes the largest enzyme cost.

### F.4 Reversible mass-action kinetics

#### F.4.1 Steady-state flux

Here we will show the full derivation of the optimal solution in the case of reversible mass-action rate laws. We start by reminding ourselves that the rate law for each reaction is given by:

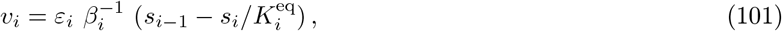

and the pathway flux is thus (as in Eq. (26)):

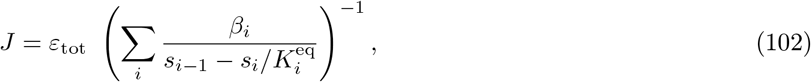

where we replaced the inequality by an equality since we are looking for the maximal flux.

While *J* can in theory be maximized by equating the gradient with respect to all metabolite levels to 0, the resulting system of equations would be difficult to solve. Instead, we will use a different approach that involves finding an expression for *J* as a function of the different enzyme levels (rather than metabolite levels).

First, we use Eq. (101) to obtain a formula for *s*_*i*_. By equating *v*_*i*_ = *J* and applying it recursively we get:

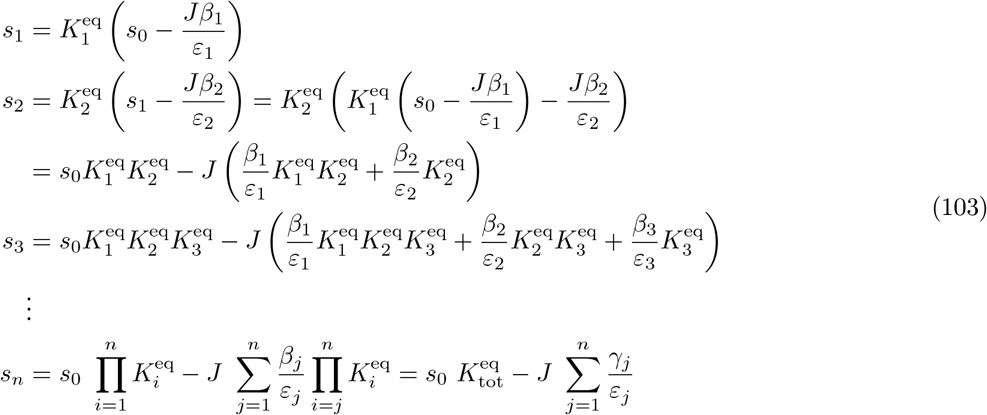

where we use the fact that 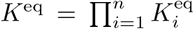 (i.e. the equilibrium constant of the pathway net reaction), and substituting 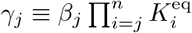. Solving for *J* we get:

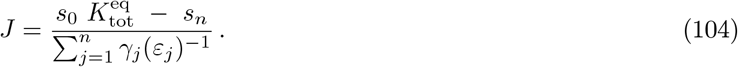

#### F.4.2 Flux control coefficients

The steady-state flux is given by Eq. (104). By taking the partial derivative with respect to enzyme levels, we obtain the flux response coefficients

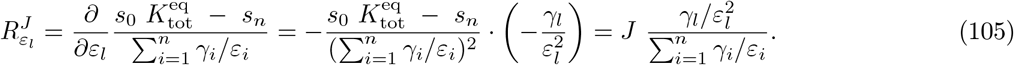

After dividing by the enzyme elasticities 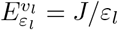, we obtain the flux control coefficients

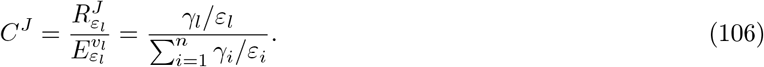

#### F.4.3 Minimum enzyme demand / maximal flux

Now, in order to maximize *J* we need to minimize the denominator (the numerator is a constant). The variables (*ε*_*j*_) have to satisfy the constraint Σ_*j*_ *ε*_*j*_ *≤ε*_tot_. To find the minimal value for the denominator, we use the Lagrange method:

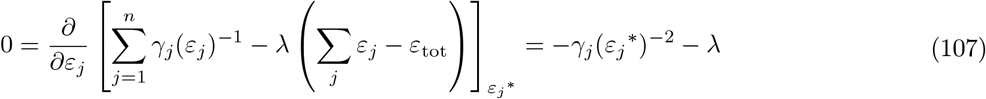

which shows that 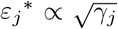. Since we know the sum of all the enzyme levels, *ε*_tot_, we can also find the proper scaling factor, i.e.:

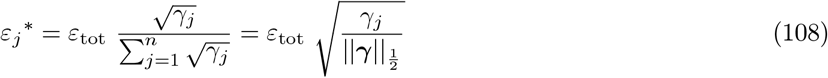

where ***γ*** is the vector of all *γ*_*j*_, and 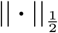 is the l_1*/*2_ norm.

Furthermore, we can calculate the maximal achievable flux *J* ^*∗*^ by plugging in the *ε*_*j*_^*∗*^ values:

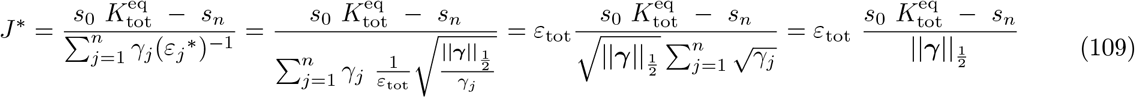

An interesting consequence of Eq. (108) is that we can draw this optimization, is looking at optimal ratios between consecutive enzymes:

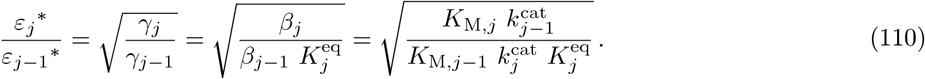

Interestingly, the ratio depends only on the *K*_M_ and *k*^cat^ values of the two reactions and is independent of all other enzymes in the system. This might be a design principle supporting for the existence of regulatory mechanisms that ensure these ratios are maintained even if the total expression level changes.

#### F.4.4 Connectivity theorem for flux control coefficients

The connectivity theorem for flux control coefficients reads 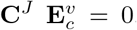. Here we verify this explicitly for an unbranched pathway with reversible mass-action kinetics. With the control coefficients Eq. (106) and the metabolite elasticities for the reversible mass action rate law

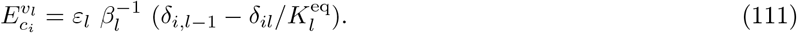

we obtain

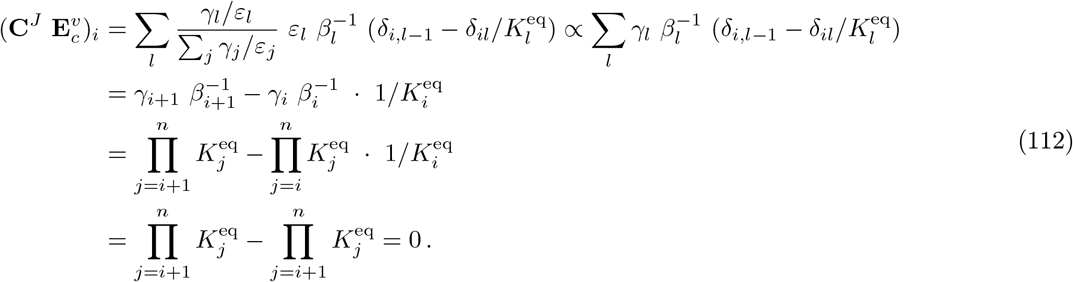

#### F.4.5 Enzyme-control rule

The enzyme levels in optimal state Eq. (27) read

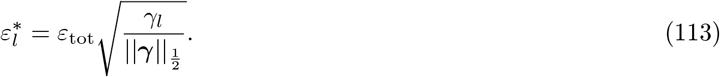

By inserting this into the formula Eq. (106) for control coefficients, we obtain (where the constant denominator is a normalisation term)

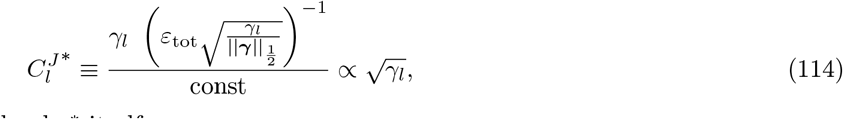

which is proportional to the enzyme level 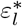 itself.

### F.5 Haldane rate law

#### F.5.1 Steady-state concentrations

For this derivation, we first use the notation in the original publication by Heinrich and Klipp [17]. For a unbranched pathway at steady-state (with flux *J* in all reactions), the Haldane rate law dictates that:

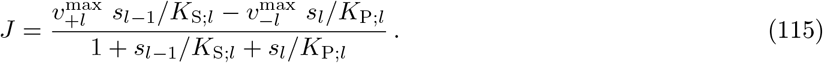

Using this formula, we can solve for *s*_*l*_ given all other parameters (including *s*_*l−*1_) and get the following recursion:

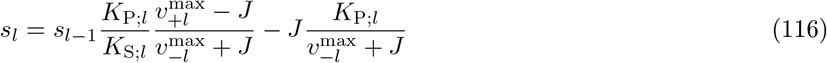

which can be solved to give:

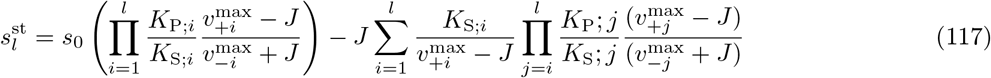

From now on, we will replace 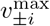 with 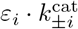 in order to better match the notation of this paper. If we use the solution for the steady-state concentration of *s*_*n*_ we can write:

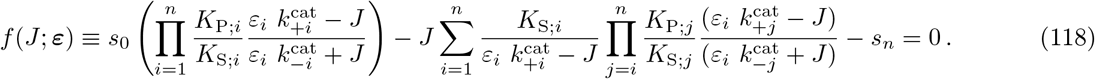

Solving this equality (i.e., finding the roots of *f* (*J* ; ***ε***)) would give us an expression for *J* (***ε***) – the flux as a function of the enzyme levels. This can be used in different ways. First, we can determine the flux response coefficients by implicit differentiation (Lemma F.2: 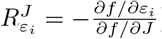), and by dividing by *J/ε*_*i*_ we obtain the flux control coefficients. Second, we can find an optimal metabolic state by minimizing Σ_*i*_ *ε*_*i*_ at a given flux *J*, and requiring Eq. (118) as a constraint (with Lagrange multiplier *λ*). Optimization using the Lagrange method would lead to the optimality condition *∂f/∂ε*_*i*_ = *λ*. However, since the derivatives are complicated and *J* and *λ* are unknown, this is hard to solve.

The most efficient method we currently have for solving this optimization problem is, unfortunately, not analytical. First, we start by writing *ε*_tot_*/J* as a function of the metabolite concentrations:

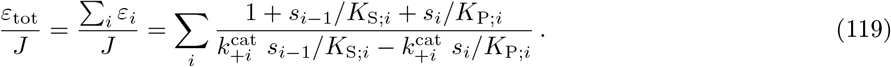

We showed previously that this function is convex with respect to the metabolite log-concentrations (ln **s**) and therefore has a single (global) minimum which is easy to find numerically [38, 8].

### F.6 Reactions with cofactors, side substrates, and side products: effective kinetic constants

The pathways considered in this paper consist of uni-uni reactions S ↔P. However, all the results apply also to reactions with several substrates and products if only one of the substrates and one of the products have variable concentrations, while all other concentrations are predefined. As an illustrating example, we consider a reaction with stoichiometric coefficients of 1, substrates *s* (variable) and *a* (constant), and products *p* (variable) and *b* (constant). There are various ways to generalize the Haldane rate law to reactions with several substrates or products. One of them, the convenience kinetics [34], for this reaction reads

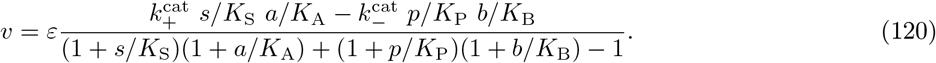

With the abbreviations *σ* = *a/K*_A_ and *π* = *b/K*_B_, we obtain

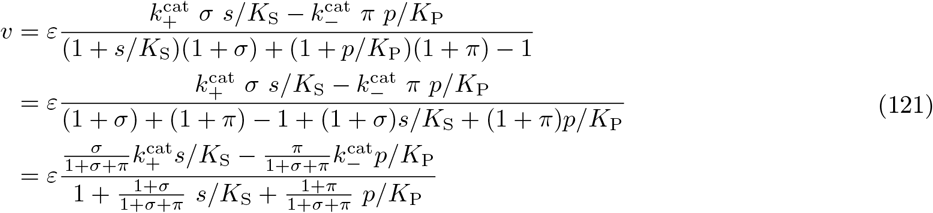

and by defining

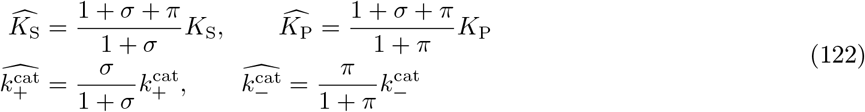

we can write this in the form of our uni-uni rate law:

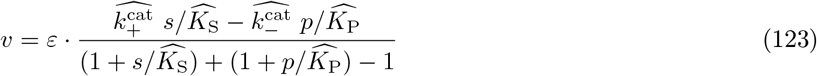

with effective 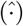 parameters. From the Haldane relationship, we obtain the effective equilibrium constant:

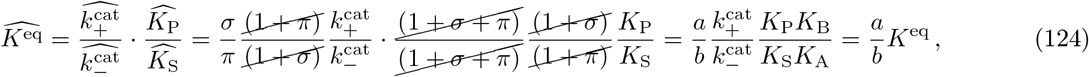

where we used the Haldane relationship for the original reaction: 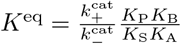.

The driving force 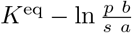 of a reaction, as a thermodynamic quantity, cannot depend on how the rate law is written. With the original rate law, it depends on the (true) equilibrium constant as well as on all four reactant concentrations. But if we ignore A and B in the rate law and simply define our equilibrium constant as the ratio *b*^*eq*^*/a*^*eq*^ in some imagined equilibrium state, the resulting driving force would be different from our original driving force and the term 1 −e_^*−θ*^_ will be wrong or, even worse, the new *θ* will have the opposite sign and be opposite to the flux direction. To avoid this problem, there are two possibilities: either we use the effective equilibrium constant Eq. (124), which contains the constant ignored concentrations, or we compute the driving force with an “external” extra term,

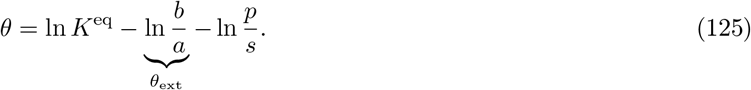

If we compare the thermodynamic force to voltage in an electric circuit, A and B would act like an external voltage source.

### F.7 Enzyme-control rule in models with constraints on enzyme and metabolite levels

We assume an unbranched metabolic pathway in steady state. The steady-state fluxes **v** = **v**^st^(***ε***) and steady-state metabolite concentrations **s** = **s**^st^(***ε***), differentiated by *ε*_*l*_, yield the response coefficients 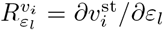 and 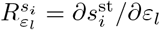. We now consider the optimality problem (same as in Eq. (40)):

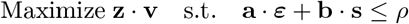

where **v, s**, and ***ε*** denote reaction fluxes, metabolite concentrations, and enzyme levels. The maximization objective is a linear benefit function scoring the fluxes, **z · v** with linear weights *z*_*i*_, while the constrained density is linear in all the concentrations, with linear weights *a*_*l*_ (for enzymes) and *b*_*i*_ (for metabolites), e.g. representing effective molecule sizes. Optimization with a Lagrange multiplier *λ* yields

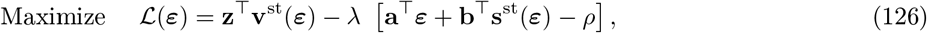

where **z** is the flux weight vector, **a** and **b** are the molecular masses of the enzymes and metabolites (respectively), and *ρ* is the upper bound on the total density. The Lagrangian yields the optimality condition:

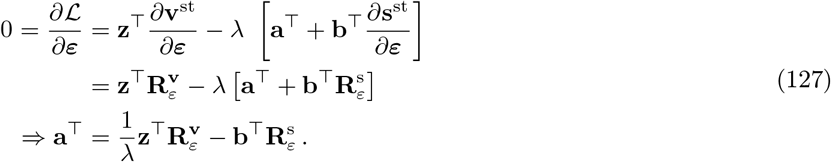

By splitting the response matrices into control and enzyme elasticity matrices, 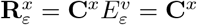diag(**v**) diag(***ε***)^*−*1^, we next obtain

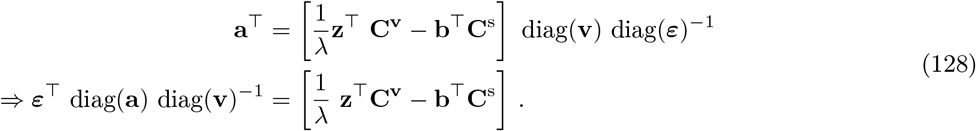

In the case of an unbranched pathway, i.e. with the same steady-state 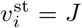 in all reactions, we can arbitrarily chose a benefit function **z**^*⊤*^ = (1, 0, …, 0) which will give use **z**^*⊤*^**v** = *v*_1_ = *J*. In this case **z**^*⊤*^**C**^**v**^ will simply be the first column of the **C**^**v**^ matrix, which we will refer to as **C**^*J*^. In addition, we can choose density constraints with uniform weights for all enzymes and for all metabolites (**a** = *a***1, b** = *b***1**). Then, the enzyme profile becomes:

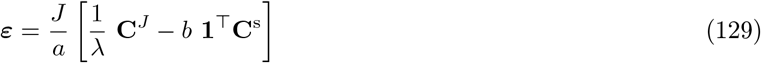

Our new enzyme-control rule thus reads, as a proportionality (and marking again the optimal state by ^*∗*^)

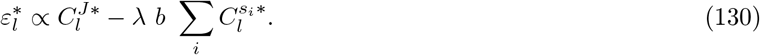

In our density constraint in Eq. (40), with a predefined overall density *ρ*, there is no explicit bound *ε*_tot_ on the sum of enzyme levels. However, given the sum of enzyme levels 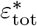 that emerges from the solution, we obtain an explicit enzyme-control rule with no unknown parameters. To derive it, we write Eq. (129) in the form

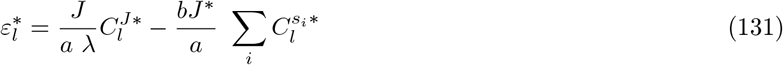

We can then sum over *l* and get:

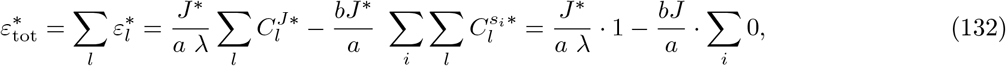

where in the last step we used the summation theorems for fluxes and metabolite concentrations. We can solve this for *λ* = *J* ^*∗*^ (*a ε*_tot_)^*−*1^, and the optimal enzyme levels read

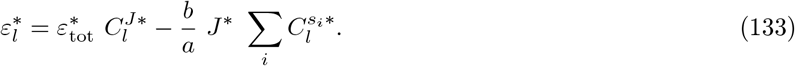

### F.8 Sufficient condition for stable states in unbranched pathways

Here we show, that a steady state in an unbranched metabolic pathway is stable if (but not only if) the first reaction 1 is reversible (i.e. it has a non-zero product elasticity), if the last reaction is not completely saturated (i.e. it has a non-zero substrate elasticity), and if in all other reactions the substrate elasticity is larger than the (absolute) product elasticity. Note that for reversible mass-action rate laws, the first two conditions are automatically satisfied and the latter condition is satisfied if *k*_+*i*_ *> k*_*−*(*i−*1)_.

**Proof** A state is asymptotically stable if all the Jacobian eigenvalues are negative (i.e. they have negative real parts). A sufficient (but not necessary) condition for stable metabolic states can be obtained from the Gershgorin’s disc theorem, which we recall here:

**Theorem F.3**. *Let* **A** *be a square matrix. Each diagonal element a*_*ii*_ *is associated with a closed disc s*_*i*_ = *D*_*i*_(*a*_*ii*_, Σ_*j*_ |*a*_*ij*_|)) *in the complex plane, with center a*_*ii*_ *and radius* Σ_*j*_ |*a*_*ij*_|. *According to the theorem, all eigenvalues of* **A** *must lie in the union of all these discs. Since the theorem applies both to* **A** *and to its transpose, each diagonal element gives rise to two discs with the same center, but different radii* Σ_*j*_ |*a*_*ij*_| *and* Σ_*j*_ |*a*_*ji*_|.

For a real-valued matrix, this means: our system is stable if for all *i, a*_*ii*_+ Σ_*j*_ |*a*_*ij*_| *<* 0 or if for all *i, a*_*ii*_+ Σ_*j*_ |*a*_*ji*_| *<* 0. We now apply this theorem to an unbranched metabolic chain with 4 reactions and 3 internal metabolites A, B, C:

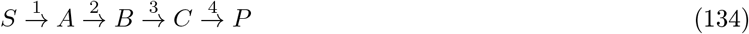

The system has no conservation relations. From the stoichiometric matrix

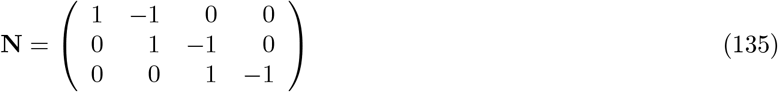

and the metabolite elasticity matrix

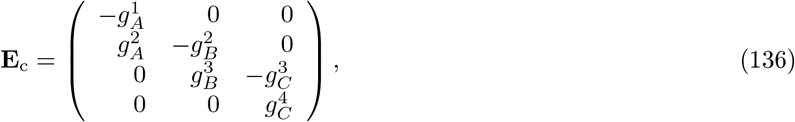

parameterised by positive values *g*, we obtain the Jacobian matrix

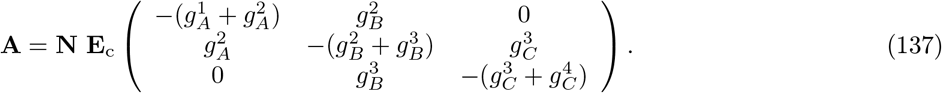

Applying Gershgorin’s theorem, from the condition for matrix columns^5^, we obtain the sufficient stability conditions

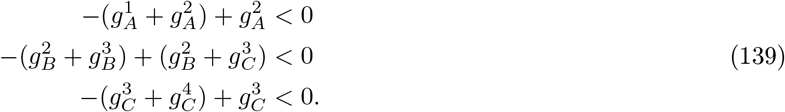

To guarantee a stable steady state, all three inequalities must hold. They are satisfied whenever 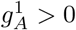 (reaction 1 is not completely irreversible), 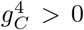 (reaction 4 is not completely saturated), and 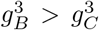 (the substrate elasticity of reaction 3 is larger than the (absolute) product elasticity of this reaction). For longer metabolic chains, we obtain the same type of conditions: the steady state will be stable if (but not only if) the first reaction is not completely irreversible, the last reaction is not completely saturated, and in every reaction in between, the substrate elasticity is larger than the (absolute) product elasticity.

### Footnotes

1 To derive this rule, we note that in an optimal state all enzymes have a marginal cost (contribution to the enzyme budget, per mol of enzyme) which must be balanced by the same marginal benefit (contribution to the production flux, per mol of enzyme). If all enzymes contribute to the enzyme budget with equal weights, their marginal costs are the same. This means that also their marginal benefit must be the same, which are given by the (unscaled) flux response coefficients 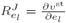. This, finally, means that their flux control coefficients 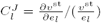 must be proportional to the enzyme levels *e*.

2 In fact, there are a number of similar arguments why the enzyme-control rules fails in this case: first, the state with infinite concentrations does not exist mathematically, so formally we cannot even refer to it as a metabolic state; second, even if this state existed, all elasticities would vanish, and the Jacobian matrix would not be invertible, so the control coefficient would not be defined; third, even if we argued that, logically, the first enzyme must have full flux control, the same argument would also apply to all other enzymes: each of the enzymes would have full flux control, thus violating the summation theorem.

3 Instead of a linear flux objective, a nonlinear function *z*(**v**) could be used. In this case, the rule (42) would remain the same, but with the gradient ∇_*v*_*z*(**v**) replacing the weight vector **z**.. Likewise, in a model with separate density constraints **a ·*ε ≤****ρ*_*ε*_ and **b ·s ≤***ρ*_*c*_, we obtain a similar formulae, but with separate Lagrange multipliers for the two constraints.

4 For a simple constraint on the total mass, the weights are simply molecular weights. This could be a proxy for excluded volume (which would still ignore, for example, hydration shells). However, a “density constraint” need not represent space demands; it may also be related to osmotic effects or opportunity costs (e.g. energy demand for production of compounds in growing cells). Therefore the meaning of the weights in the density constraint may differ from model to model.

5 Incidentally, from the condition for rows we obtain another (alternative) set of sufficient conditions 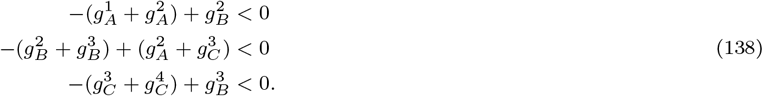 These conditions are satisfied, for example, if all elasticities are non-zero, if the product elasticity 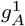 of the first reaction is equal or larger than the product elasticity 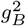 of the second reaction, if the substrate elasticity of the last reaction is equal or larger that the substrate elasticity 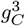 of the third reaction, and if 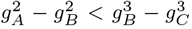. For a reversible mass-action kinetics *v* = *e*[*k*_+_*s − k*_*−*_*p*] the latter condition would mean: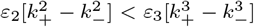 or 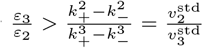 where the “standard velocity” 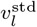 denotes the rate that a reaction *l* would show at unit metabolite and enzyme levels.

